# Structural Basis for the Clamping and Ca^2+^ Activation of SNARE-mediated Fusion by Synaptotagmin

**DOI:** 10.1101/574632

**Authors:** Kirill Grushin, Jing Wang, Jeff Coleman, James E. Rothman, Charles E. Sindelar, Shyam S. Krishnakumar

**Affiliations:** Department of Cell Biology, Yale University School of Medicine, 333 Cedar Street, New Haven, CT 06520, USA; Department of Molecular Biophysics and Biochemistry, Yale University School of Medicine, 333 Cedar Street, New Haven, CT 06520, USA.; Department of Clinical and Experimental Epilepsy, Institute of Neurology, University College London, Queens Square House, London WC1 3BG, UK.

**Keywords:** Synaptotagmin, SNARE, cryo-electron microscopy, calcium, lipid nanotubes

## Abstract

Synapotagmin-1 (Syt1) interacts with both SNARE proteins and lipid membranes to synchronize neurotransmitter release to Ca^2+^-influx. To understand the underlying molecular mechanism, we determined the structure of the Syt1-SNARE complex on lipid membranes using cryo-electron microscopy. Under resting conditions, the Syt1 C2 domains adopt a novel membrane orientation with a Mg^2+^-mediated partial insertion of the aliphatic loops, alongside weak interactions with the anionic lipid headgroups. The C2B domain concurrently binds the SNARE bundle via the ‘primary’ interface and is positioned between the SNAREpins and the membrane. In this configuration, Syt1 is projected to sterically delay the complete assembly of the associated SNAREpins and thus, contribute to clamping fusion. This Syt1-SNARE organization is disrupted upon Ca^2+^-influx as Syt1 reorients into the membrane, allowing the attached SNAREpins to complete zippering and drive fusion. Overall, we find cation (Mg^2+^/Ca^2+^) dependent membrane interaction is a key determinant of the dual clamp/activator function of Syt1.

---

The rapid Ca^2+^-triggered release of neurotransmitters at the synapse is a highly orchestrated process^1-3^. The proteins involved are known and are well-characterized^1-4^. This includes the SNARE (Soluble N-ethylmaleimide-sensitive factor attachment protein receptor) proteins (VAMP2, Syntaxin, and SNAP25) that catalyze synaptic vesicle (SV) fusion as well as chaperones Munc13, Munc18 and regulators Complexin and Synaptotagmin^1-4^. Synaptotagmin-1 (Syt1) is a key component involved in all stages of the process, including SV docking and priming ^5-9^, preventing un-initiated SV fusion^10,11^ and triggering fusion upon Ca^2+^ influx^12-15^. Remarkably, such broad specialization has not resulted in structural complexity of the protein.

Syt-1 is anchored in the SV by a transmembrane domain (TMD), which is connected to tandem cytosolic C2 domains (C2A & C2B) via a 60-residue flexible linker. Both C2 domains are eight stranded anti-parallel beta sandwiches with one calcium sensing region consisting of two aliphatic loops containing Aspartic acid (Asp) residues that bind Ca^2+^^16-18^. These conserved Asp residues coordinate Ca^2+^ with the acidic lipids, like phosphatidylserine (PS) and phosphatidylinositol 4, 5-bisphosphate (PIP2), to facilitate the membrane insertion of the flanking aliphatic loops^5,18-21^. In addition, the C2B domain has a conserved poly-lysine motif that binds to PS/PIP2 headgroups to mediate Ca^2+^-independent membrane interaction^5,7,8^. Both the Ca^2+^-independent and dependent membrane interactions of the Syt1 C2B domain are functionally critical. The interaction of the poly-lysine motif with PIP2 clusters facilitates the initial docking of the SV at the active zone ^6,7,22,23^ and Ca^2+^-dependent membrane insertion of the aliphatic loops is physiologically required for triggering synaptic transmission^13,21,24,25^. The C2A domain does not have a pronounced polybasic patch like C2B and its Ca^2+^-dependent membrane interaction is not absolutely required for neurotransmission^13,26,27^. Nevertheless, recent studies show that the C2A domain can modulate the Syt1 function^28-31^.

Genetic analysis shows that Syt1 also plays a critical role in ‘clamping’ spontaneous release and Ca^2+^-evoked asynchronous release^11,15,32^. Similarly, the deletion of Complexin (Cpx) shows increase in spontaneous release frequency and de-synchronization of evoked response^33-35^. This suggests that Syt1 and Cpx act ‘synergistically’ to clamp un-initiated/delayed release components to ensure Ca^2+^-coupled synchronous transmitter release. Under *in vitro* conditions, Syt1 and Cpx have been shown to independently clamp SNARE-mediated vesicle fusion under specific conditions^36-38^, but both Syt1 and Cpx are required to constitute a Ca^2+^-dependent clamp under physiologically-relevant conditions^39,40^. However, the precise molecular mechanism of clamping is still unclear.

Besides membranes, Syt1 also interacts with the SNARE proteins^22,23,41,42^. Specifically, it binds to the Syntaxin/SNAP25 complex (t-SNAREs) on the plasma membrane (PM). This interaction is crucial for Syt1 function at all stages of SV exocytosis^22,23,41,42^. Crystal structures of Syt1-SNARE complexes in the presence and the absence of Cpx^41,42^ revealed two main interaction sites for the C2B domain on the t-SNARE protein. One site (“primary”) is Cpx-independent and involves both helices of SNAP-25. The second site is Cpx-dependent (“tripartite”) and involves the portions of helices of Cpx and Syntaxin^41,42^. Both of these sites were shown to be important for Syt1 clamping and Ca^2+^-activation function^41,42^. However, it is still unclear if and how these interactions manifest under the membrane environment. Recent efforts to visualize the protein organization at the contact sites of reconstituted vesicles using cryo-electron tomography generated structures which does not have sufficient resolution to readily distinguish the interaction partners or their organization^43^.

Based on the currently available structural, biochemical and physiological data, several speculative models for Ca^2+^-regulated SV exocytosis have been proposed^44-48^. But the precise molecular architecture that accommodates all the functionally relevant SNARE and membrane interactions of Syt1 is unclear. This information would be crucial towards understanding the protein organization of the ‘clamped’ state and the underlying mechanisms that enable rapid and Ca^2+^-synchronized neurotransmitter release.

In this study, we sought to acquire structural information of the Syt1-SNARE interaction in the presence of a negatively-charged lipid membrane. Using cryo-electron microscopy (Cryo-EM), we obtained 10Å resolution reconstruction of the Syt1-SNARE complex on lipid membranes in the presence of Mg^2+^. This structure, derived from a helical crystal formed by covalently linked Syt1-SNARE protein applied to lipid tube surface, revealed that the Syt1 C2B domain concurrently binds both the SNARE proteins and the negatively charged membranes on diametrically opposite surfaces. It showed that the Syt1 C2B domain interacts with the SNARE complex via the recently described ‘primary’ interface on the t-SNAREs. At the same time, both Syt1 C2 domains adopt a previously unknown orientation with one of the aliphatic loops (loop 3) partially submerged into the phospholipid headgroups. This arrangement suggests a straightforward clamping mechanism as Syt1 is ideally positioned to create a steric block hindering the full assembly of the associated SNAREpins. In contrast, the Syt1-SNARE complex failed to organize on the lipid surface in the presence of Ca^2+^. The overall protein appearance suggests that upon Ca^2+^-binding, the Syt1 C2 domains reorient into the membrane, likely displacing the associated SNAREpins from the primary interface. Our data indicates that the Ca^2+^ trigger of SV fusion possibly involves large conformational changes in the associated proteins.

## RESULTS

### Syt1^C2AB^-SNARE Complex Helically Organizes on Lipid Tubules

Our attempts to co-crystallize Syt1 and SNARE on membranes using individual proteins were unsuccessful. So, we used a protein construct wherein the minimal C2AB domains of Syt1 (Syt1^C2AB^) are covalently linked to the SNARE complex, which was proven to be successful in 3D crystal formation^41^. Negative stain EM analysis of Syt1^C2AB^-SNARE protein complex incubated with liposomes containing negatively charged lipids (DOPC/DOPS/PIP2 60/34/6) revealed the formation of protein-coated lipid tubules with helical diffraction peaks (Supplementary Fig. 1). These protein-coated tubular projections, formed in the presence of 1 mM free Mg^2+^, occurred in low frequency and were quite variable in size, with diameters ranging between ∼40 nm and ∼60 nm (Supplementary Fig. 1). Cryo-EM analysis further showed that the lipid tubules have inconsistent protein decoration, wherein some were partially decorated with protein, while other tubules were devoid of protein (Supplementary Fig. 1).

To improve the quantity and quality of the protein coated lipid tubes suitable for the helical 3D reconstruction process, we applied Syt1^C2AB^-SNARE protein to pre-formed lipid nanotubes (LNT) (Supplementary Fig. 2). The LNTs of fairly uniform size (diameter ∼35 nm) were assembled with the inclusion of glycolipid (20% galactosylceramide, GalCer 24:1) into the vesicle lipid mixture^49^. Glycolipid derived nanotubes are routinely employed as substrate for preparing 2D-crystals, as they promote the formation of helical arrays of adsorbed proteins and macromolecular assemblies^50^. For example, they were recently used to helically organize and reconstruct the membrane bound blood coagulation factor VIII containing C2 domains by Cryo-EM^51^.

Correspondingly, negative stain EM analysis showed a large number of homogeneous protein coated LNTs, with a strong diffraction pattern corresponding to a helical organization of protein molecules on the lipid surface. Notably, these protein-coated LNTs, formed in presence of 1mM free Mg^2+^, were analogous to the tubules derived from vesicles both in their appearance and diffraction peaks (Supplementary Fig. 2). But frozen LNT samples analyzed by CryoEM revealed a majority of unorganized or poorly diffracted tubes (Supplementary Fig. 2). Since the partial protein decoration was never observed by negative stain EM for the vesicle-derived or pre-assembled tubules, we reasoned that the freezing likely weakens the Syt1-membrane interaction and the proteins were falling off the tube surface during the Cryo-EM grid preparation. So, to ensure tight binding of the proteins on the surface and to improve their packing, we increased the acidic lipid content and simplified the lipid composition of the LNTs. The optimum composition was found to be DOPS: GalCer at 80:20 ratio, which vastly improved both the overall frequency and quality of the helically organized Syt1^C2AB^-SNARE coated tubes (Fig. 1A). This enabled us to collect a homogenous dataset suitable for high-resolution reconstruction of the Syt1-SNARE complex bound to membranes under resting conditions (1mM Mg^2+^).

**Figure 1.**
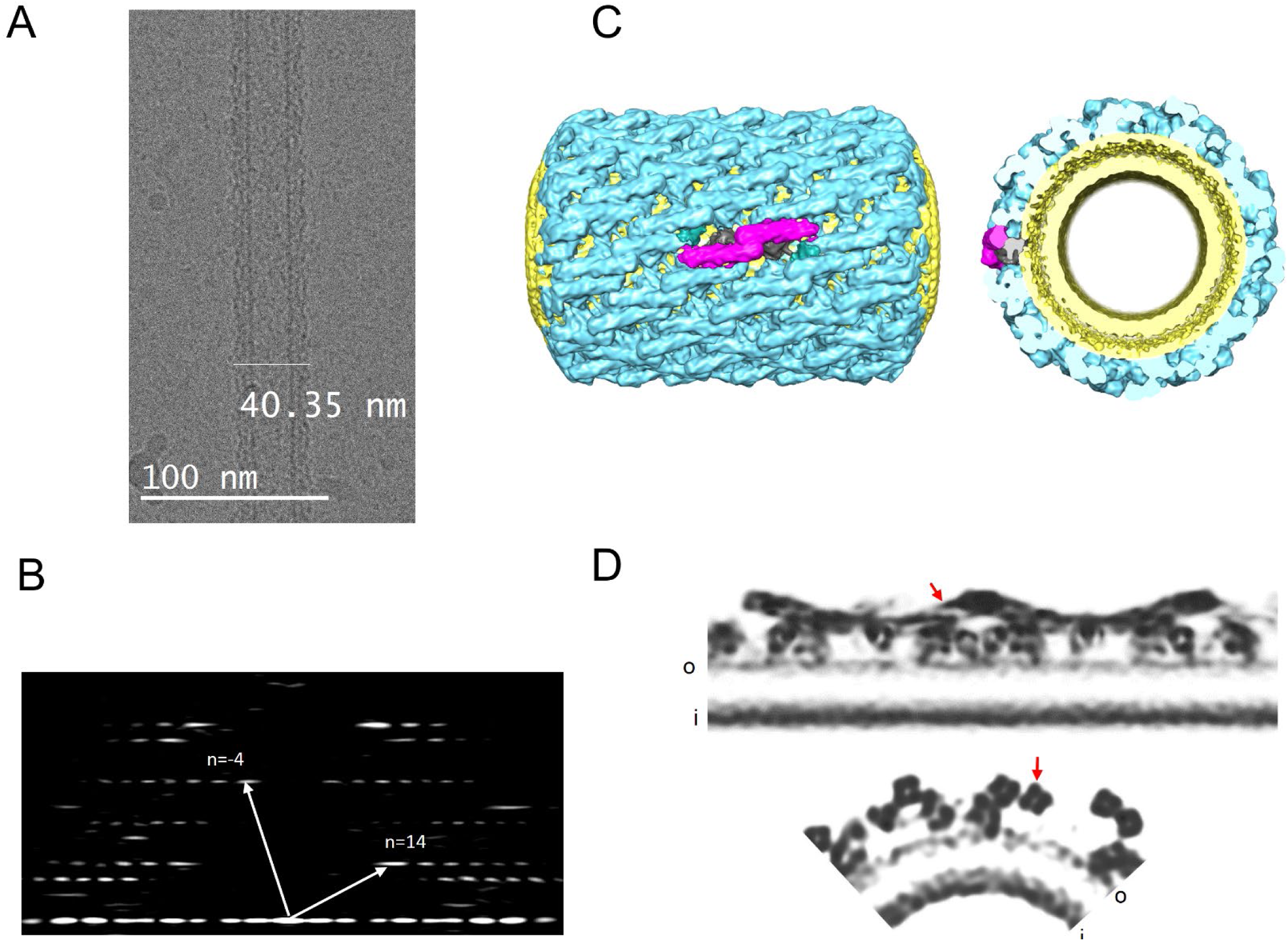
Helical crystallization of the Syt1^C2AB^-SNARE complex on the lipid membrane surface. (A) Cryo-electron micrograph of the Syt1^C2AB^-SNARE complex helically organized on lipid nanotubes. A representative micrograph of ∼41 nm protein-coated LNTs with homogenous helical packing used in the reconstruction is shown. (B) Corresponding Fourier Transform of the LNTs shows a helical diffraction pattern with the Bessel functions for basis vectors, which corresponds to Δz=7.35 A Δφ=78.48; 14,-4 start helix. (C) Iso-surface representation of the helical reconstruction of protein-coated LNTs at 10.4 Å resolution. The Syt1^C2AB^-SNARE protein (blue) is arranged on the lipid bilayer tube (yellow) with characteristic rod-like densities (magenta) corresponding to the SNARE complex oriented almost parallel to the tube axis and thus, locally experience a relatively-flat bilayer surface. (D) Grayscale density slice through the one side of the 3D-volume along the helical axis (top, slice depth 1) and quarter grayscale slice from cross-sectional density (bottom, slice depth 30). The densities corresponding to a-helices forming the SNAREpin (red arrow), as well with phospholipid heads plane of the inner (i) and outer (o) membrane leaflets are also well defined.

### CryoEM Structure of Syt1^C2AB^-SNARE on Lipid Surface

We evaluated the quality of protein coating by analyzing the power spectra of selected tubes and their suitability for helical indexing by the Fourier-Bessel method^52^. Protein decorated LNTs around ∼41 nm (outer diameter) showed helical diffraction corresponding to homogenous helical crystal (Fig. 1B), so this subset of tubes was selected for further structural analysis. Helical indexing of the representative tube defined the Syt1^C2AB^-SNARE arrangement on the surface as 14,-4 start helix with a helical rise (Δz) of 7.35 Å and a helical twist (Δφ) of 78.48° (Fig. 1B). These parameters were then used for the iterative helical real-space reconstruction (IHRSR)^53^. The resulting 3D reconstruction at 10.4 Å resolution showed protein decoration on the tube surface with prominent rod-like features (SNARE proteins) arranged almost parallel to the tube axis and with globular structures (C2 domains) attached to the membrane surface (Fig. 1C & D). SNARE complexes were well resolved with distinguishable densities corresponding to the α-helices of SNARE proteins, however the orientation of SNAREs and relative positioning of the Syt1 C2A and C2B domains was not clear. Noteworthy, the proteins were arranged parallel to the nanotube axis (Fig. 1C & D) and thus, locally experience relatively flat-bilayer surfaces.

We noticed a striking visual similarity of protein organization between the 2D-crystal on the membrane and the recently published 3D crystal of the same complex^41^. So, we performed rigid body docking with separate C2A, C2B and SNAREs derived from the Syt1-SNARE crystal structure (PDB code: 5CCI^41^). The crystal structures were unambiguously docked into the EM map and showed SNAREpins arranged in an antiparallel fashion with membrane-distal C-terminal interaction. The Syt1 protein located between the SNARE complex and membrane surface (Fig. 2, Supplementary Fig. 3, Supplementary Video 1), with the C2B domain interacting with the middle portion of SNARE complex. Relative position of C2B over the SNAREs was very similar to the recently described ‘primary’ binding site, involving the t-SNARE helices^41,42^. The linker between the C2A and C2B domains was fully extended, with the C2A domain locating close to but not interacting with the N-terminal end of the same SNAREpin (Fig. 2, Supplementary Fig. 3). In parallel, we also aligned the crystal structure of a single Syt1^C2AB^-SNARE complex (primary interface unit) from 5CCI X-ray structure onto our CryoEM-derived Syt1^C2AB^-SNARE reconstruction (Supplementary Fig. 4). It showed that in relation to the SNARE complex, the Syt1 C2B domain is identically positioned and nearly superimposable, but the C2A position deviates slightly compared to the crystal structure (Supplementary Fig. 4). This indicates that on the membrane surface, the primary interface between C2B and SNARE proteins is preserved, but the C2A domain is likely more flexible. Overall, the Syt1^C2AB^-SNARE complexes were more densely organized on the lipid surface, and by conforming to the curved surface of the LNTs their higher order organization deviated from the 3D crystal lane (Supplementary Fig. 4).

**Figure 2.**
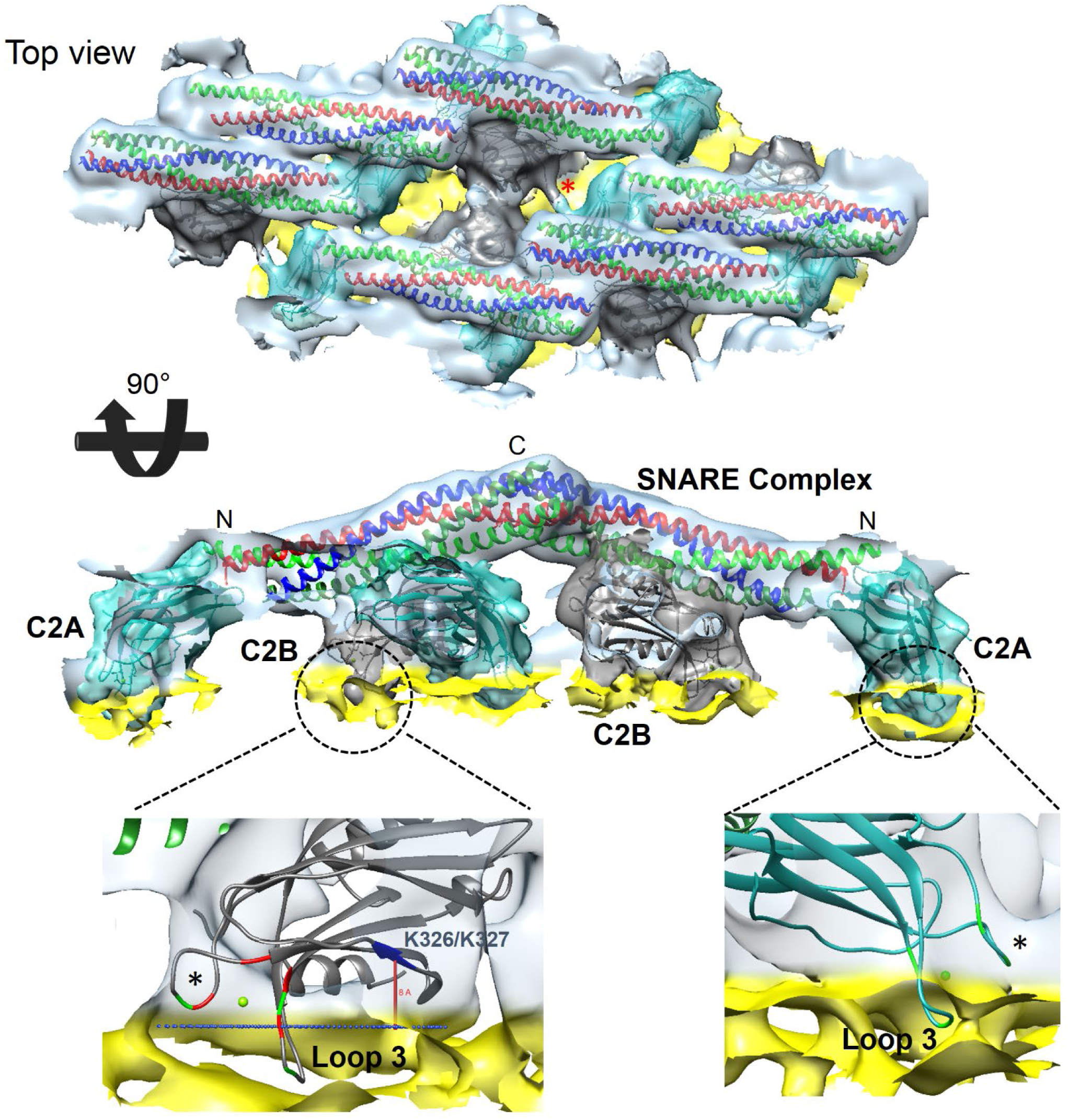
Organization of Syt1^C2AB^-SNARE on a lipid membrane surface. The rigid body docking of single C2A, C2B, SNARE crystal structures from Syt1^C2AB^-SNARE 3D crystal (5CCI) onto the Cryo-EM map. SNAREpins (multi-color) form the top-most protein layer and are positioned in antiparallel fashion with their C-terminus pointing away from the membrane (yellow). The Syt1 C2B domains (gray) are positioned between SNAREpins and the outer membrane’s phospholipid heads plane. The linker between the C2A (cyan) and C2B domain is extended and clearly visible (red asterisk) allowing unambiguous identification of C2A and C2B domains within the same Syt1^C2AB^ unit. Both C2 domains adopt a previously unseen orientation relative to the membrane surface. One of the Ca^2+^-coordinating aliphatic loops (loop 3) on both C2 domains is partially inserted into the phospholipid heads plane, while loop 1 (black asterisk) stays close to the lipid surface. The conserved polybasic patch (residue K326/327 in blue) in the C2B domain is also positioned close to the lipid head group, suggesting possible interaction.

The Syt1^C2AB^-SNARE complex adopted a previously undescribed orientation on the membrane surface with the C-terminal end of the SNAREs pointing away from the membrane and the Syt1 C2A and C2B calcium loops oriented towards the membrane (Fig. 2). Fitting the C2A and C2B domain crystal structure showed that one of the Ca^2+^-coordinating loops (loop 3) of both C2 domains was submerged into the phospholipid heads plane of the outer membrane leaflet (Fig. 2, Supplementary Fig. 3). Notably, even at this partially inserted state, the conserved polylysine motif of the C2B domain, located within the expected distance range (∼8Å) from the phospholipid head group^54^. Reinforcing this reconstruction, we observed similar overall organization of the Syt1^C2AB^-SNARE complex on vesicle derived tubules and LNTs assembled with PS/PIP2, albeit the low resolution of the resulting structures (Supplementary Fig. 5). Under all conditions, both C2A and C2B domains were positioned similarly on the membrane. Nonetheless, we chose to focus on the C2B-membrane interaction for further analysis, as its physiological relevancy is well-established compared to the C2A domain.

### Mg^2+^ Orchestrates Membrane Interaction of Syt1^C2AB^

To gain further insight into the novel membrane orientation of the Syt1^C2AB^-SNARE complex, we investigated the consequence of targeted mutations altering the membrane interaction of the Syt1 C2B domain. Specifically, we assessed its ability to form an organized structure on the LNT surface based on visual appearance and helical diffraction pattern (Fig. 3). Neutralizing the conserved polybasic motif (K326A/K327A) of the C2B domain^5^ did not significantly affect the helical organization of Syt1^C2AB^-SNARE on the LNTs, which were visually identical to the wild type (Fig.3). This suggested that the interaction of the polybasic patch with acidic lipids does not solely determine the attachment of the C2B domain to the membrane surface. Consequently, we looked into the effect of altering the physiochemical properties of the Ca^2+^-loops^55,56^. Polar mutations of aliphatic residues in the Ca^2+^-loop (V304N/Y364N/I367N; Syt1^3N^), intended to decrease its hydrophobicity and hinder its membrane insertion, abrogated the helical organization even though the protein still bound to the LNTs (Fig 3). Conversely, enhancing the hydrophobicity of the Ca^2+^-loops, by neutralizing the Ca^2+^-coordinating Asp residues (D307/D363/D365A; Syt1^3A^), stabilized the organized structures even in the absence of Mg^2+^ (Fig. 3). This was striking as the wild type Syt1 protein strictly required Mg^2+^ to form organized structures on LNTs. In fact, Mg^2+^ was crucial for the wild type Syt1^C2AB^-SNARE complex to stay bound to the LNT surface during the freezing procedure. This indicated that the Mg^2+^ stabilizes the Syt1-membrane interaction under these conditions, likely by coordinating the conserved Asp residues with the acidic lipid head group. Altogether, these results show the importance of Mg^2+^-driven C2B domain orientation with partial membrane insertion of the aliphatic loops for the Syt1^C2AB^-SNARE complex helical arrangement.

**Figure 3.**
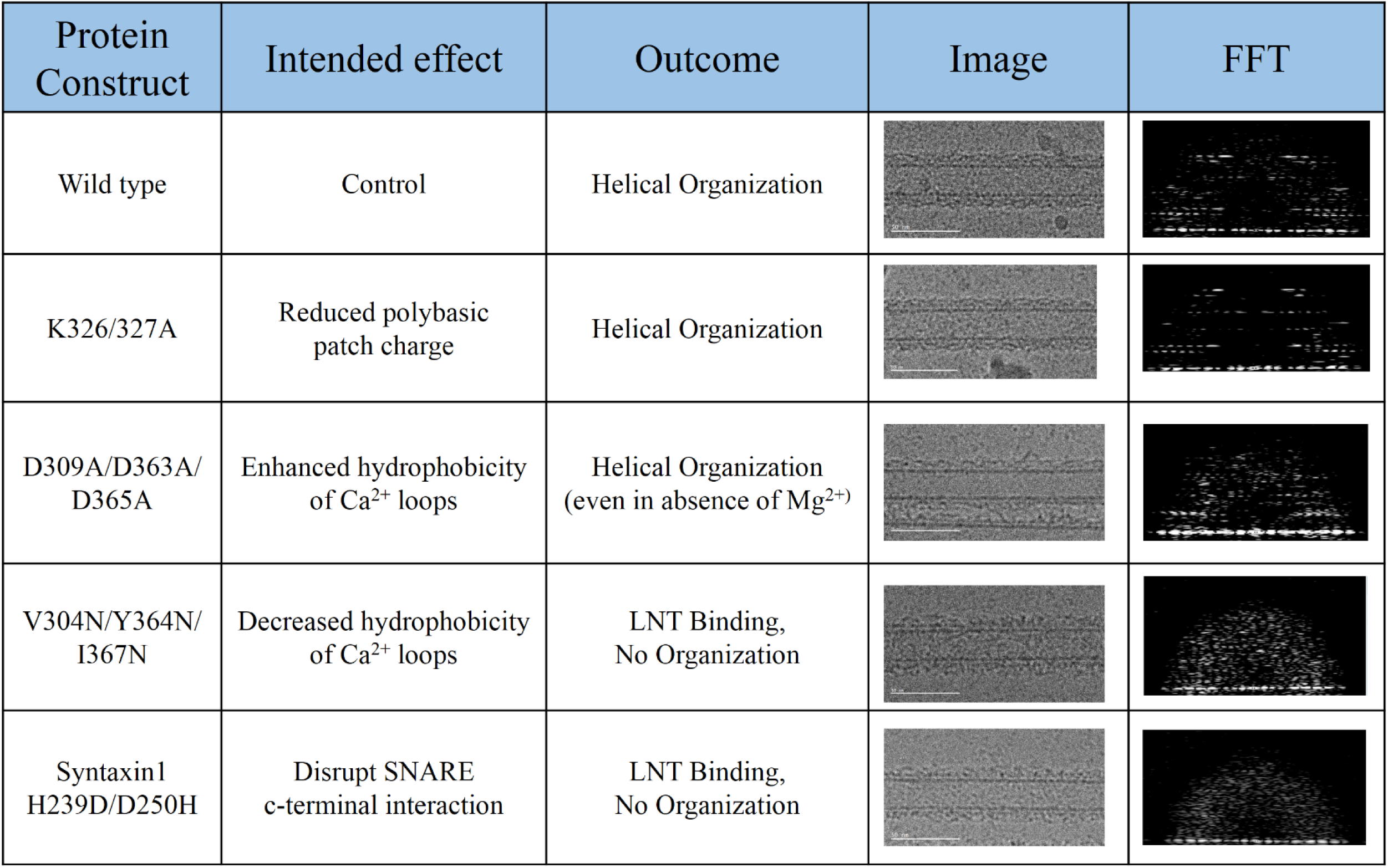
Summary of targeted mutations in the Syt1^C2AB^ and SNARE complex used to identify and validate critical crystallographic contacts. The mutations were assessed for their ability to form a helical crystal on the membrane surface by visual appearance and by helical diffraction pattern of corresponding Fourier transforms (shown as acquired at the vertical tubes orientation) of the decorated LNTs by Cryo-EM analysis. Disruption of the C2B polybasic patch (K326A/K327A) had no effect on helical organization, but alteration of the Ca^2+^-loops showed a strong effect highlighting its critical role. Disrupting the C-terminal interaction of the SNAREpin also interfered with the helical organization indicating its role in stabilizing the helical formation.

To ascertain that the partially membrane inserted state with Mg^2^ is not an artifice of crystallography, we examined if this can occur under dilute conditions using isolated proteins. We employed a well-established fluorescence assay with an environment-sensitive probe IAEDANS [5-9(((2-iodoacetyl)amino)ethyl)amino)naphthalene-1-sulfonic acid] introduced a the tip of the Ca^2+^-loops of Syt1 C2A and C2B domains to track their localization^5,20^. In the presence of negatively charged liposomes (60% PC, 34% PS, 6% PIP2), the IAEDANS probe introduced at the loop 3 of both C2A (residue 235) and C2B (residue 367) exhibited a Mg^2+^-dependent increase in fluorescence intensity (∼40% compared to the EDTA condition) and blue shift (∼15 nm) in the emission spectrum. This is consistent with IAEDANS probe locating within a hydrophobic (i.e. lipid membrane) environment (Fig. 4). In contrast, very little to no change in fluorescence signal was observed for IAEDANS probes introduced into loop 1 (residue 173 on C2A; or residue 304 on C2B) (Fig. 4, Supplementary Fig. 6). This fluorescence analysis matches perfectly with the CryoEM structure showing a partial membrane insertion of C2 domain loop 3, but not loop 1 (Fig. 2).

**Figure 4.**
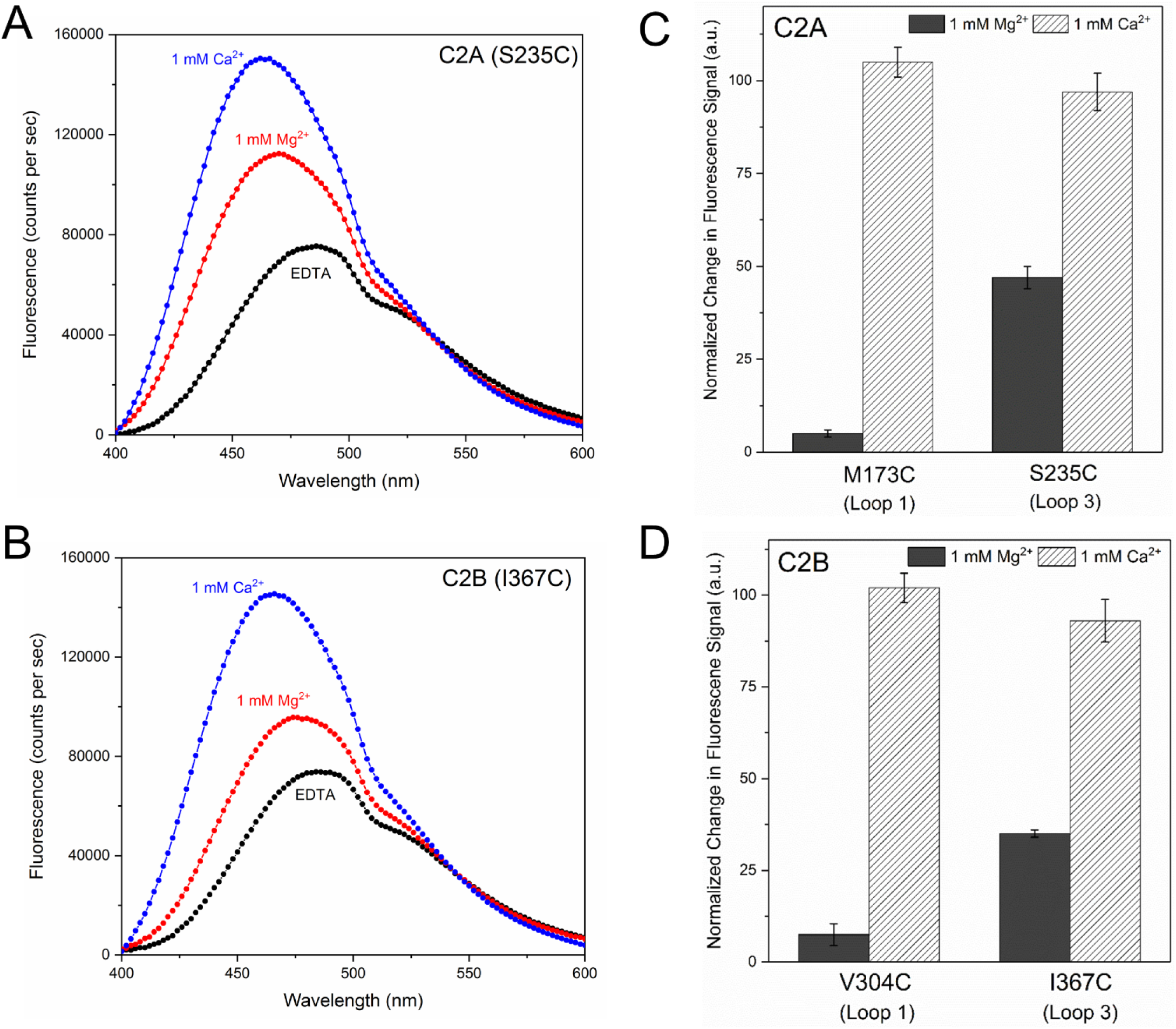
Mg^2+^ induces the partial insertion of loop 3 on both C2 domains. (A & B) Fluorescence analysis using environmentally-sensitive probe, IAEDANS introduced at the tip of the aliphatic loops of the C2 domains shows the Mg^2+^ (red curve) induces an insertion of loop 3 (residue S235 of C2A & residue I367 of C2B) as evidenced by an increase in the fluorescence intensity and blue-shift in the emission maxima as compared to an EDTA control (black curve). This Mg^2+^-induced membrane interaction is distinct, with shallower positioning of the aliphatic loops in the lipid membrane as compared to the Ca^2+^-bound state (blue curve) (C & D) Normalized fluorescence signal at the emission maxima shows that Ca^2+^-induces deep insertion of both loop 1 (residue M173 in C2A and V304 in C2B) and loop 3, but Mg^2+^ specifically triggers partial insertion of loop 3 alone in both C2 domains. Representative fluorescence curves are shown, along with averages and standard deviations from 4-5 independent experiments.

The fluorescence profiles were markedly different in the presence of 1 mM Ca^2+^. The IAEDANS probes on both loops 1 and 3 of either C2 domain exhibited a strong increase in fluorescence intensity (∼95% compared to EDTA control) and a larger blue shift (∼25 nm) in emission spectrum corresponding to stronger and deeper membrane association (Fig. 4, Supplementary Fig. 6). These results demonstrate that in the presence of 1mM Mg^2+^, loop 3 of the Syt1 C2A and C2B domains are submerged into the membrane even under non-crystallizing conditions but not as deep compared when Ca^2+^ is present. Importantly, it implies that the Mg^2+^-induced partial-membrane insertion of Syt1 is an intrinsic feature, distinct from the Ca^2+^-bound state, even though it involves the same structural elements. This is consistent with recent surface forces apparatus measurements^57^ showing that Mg^2+^ increases the binding energy of Syt1 to anionic membranes as compared to EGTA, but is weaker than the Ca^2+^-bound state (∼6 k_B_T for EGTA, ∼10 k_B_T with Mg^2+^, ∼18 k_B_T with Ca^2+^).

So, it emerges that the Mg^2+^-driven Syt1 orientation determines the relative positioning of the SNARE complex on the lipid membrane. As such, the SNARE complex bound to the Syt1 C2B domain via the primary interface is oriented with its C-terminus pointed away from the membrane surface (Fig. 2, Supplementary Fig. 3). If this Syt1-SNARE arrangement were to form under physiological conditions, it might sterically hinder the complete assembly of the membrane-anchored SNAREs and prevent fusion under resting (1mM Mg^2+^) conditions (see discussion).

In our helical structure (similar to the 3D crystal), we also observed an interaction between neighboring SNAREpins mediated by charged residues in the Syntaxin1 and VAMP2 C-termini (Supplementary Fig. 3). Disrupting this interaction, using a mutation on Syntaxin (H239D/ D250H), resulted in complete loss of helical organization on LNTs (Fig. 3), highlighting its equal importance in the helical arrangement. IAEDANS-based fluorescence experiments, including analysis of Syt1 3A/3N proteins, showed that helically organized structures are stably formed only under conditions wherein Syt1 adopts a partially-inserted state (Supplementary Figure 7). Hence, we reason that the Syt1 C2B guides the SNAREpin orientation, and the C-terminal SNARE-SNARE interaction in turn stabilizes the membrane inserted state of the Syt1 C2 domains and the helical arrangement on the membrane surface. Correspondingly, we were unable to crystalize the Syt1-SNARE complex with truncated SNARE complexes.

### Ca^2+^ Disrupts Syt1^C2AB^-SNARE Organization on the Membrane Surface

The Syt1^C2AB^-SNARE complex did not form any organized structures on LNTs in the presence of 1mM free Ca^2+^ (Fig. 5). Instead, we observed repeating densities on the membrane surface (red arrows in Fig. 5) with unorganized auxiliary densities (blue arrows in Fig. 5) on the top. Given the dimensions of the observed densities and known physiology^5,19,20^, we infer that the dense packing on the membrane (with smeared diffraction peaks) correspond to the Syt1^C2AB^ protein, while the SNARE complex, displaced from its primary binding site, represents the surrounding density. This is in stark contrast to 3D X-ray crystallography results, which shows similar Syt1^C2AB^-SNARE organization in the Ca^2+^-bound state^41^. Thus, we conclude that the Ca^2+^ induced reorientation of Syt1 into lipid membranes^5,19,20^, disrupts the Syt1-SNARE organization found under the resting conditions. Furthermore, it appears that the deep membrane insertion of the C2 domain aliphatic loops^19,20^ likely dislodges the associated SNAREpins from the primary interface. Consistent with this premise, the Syt1^3A^ protein that lacks the ability to bind Ca^2+^ and thus, insert deep into the membranes (Supplementary Figure 7) helically organized on LNTs even in the presence of Ca^2+^ (Supplementary Fig. 8). Also, Ca^2+^ restored the ability of the Syt1^3N^ mutant to organize on LNTs (Supplementary Fig. 8). Noteworthy, IAEDANS fluorescence experiment indicates that Ca^2+^ induces partial insertion of the Syt1^3N^ aliphatic loops mimicking the Mg^2+^-bound state of the wild type Syt1 (Supplementary Figure 7). In sum, we find that Ca^2+^-binding induces a large-scale conformational rearrangement of the Syt1-SNARE complex on the lipid membrane surface, disrupting the pre-fusion architecture.

**Figure 5.**
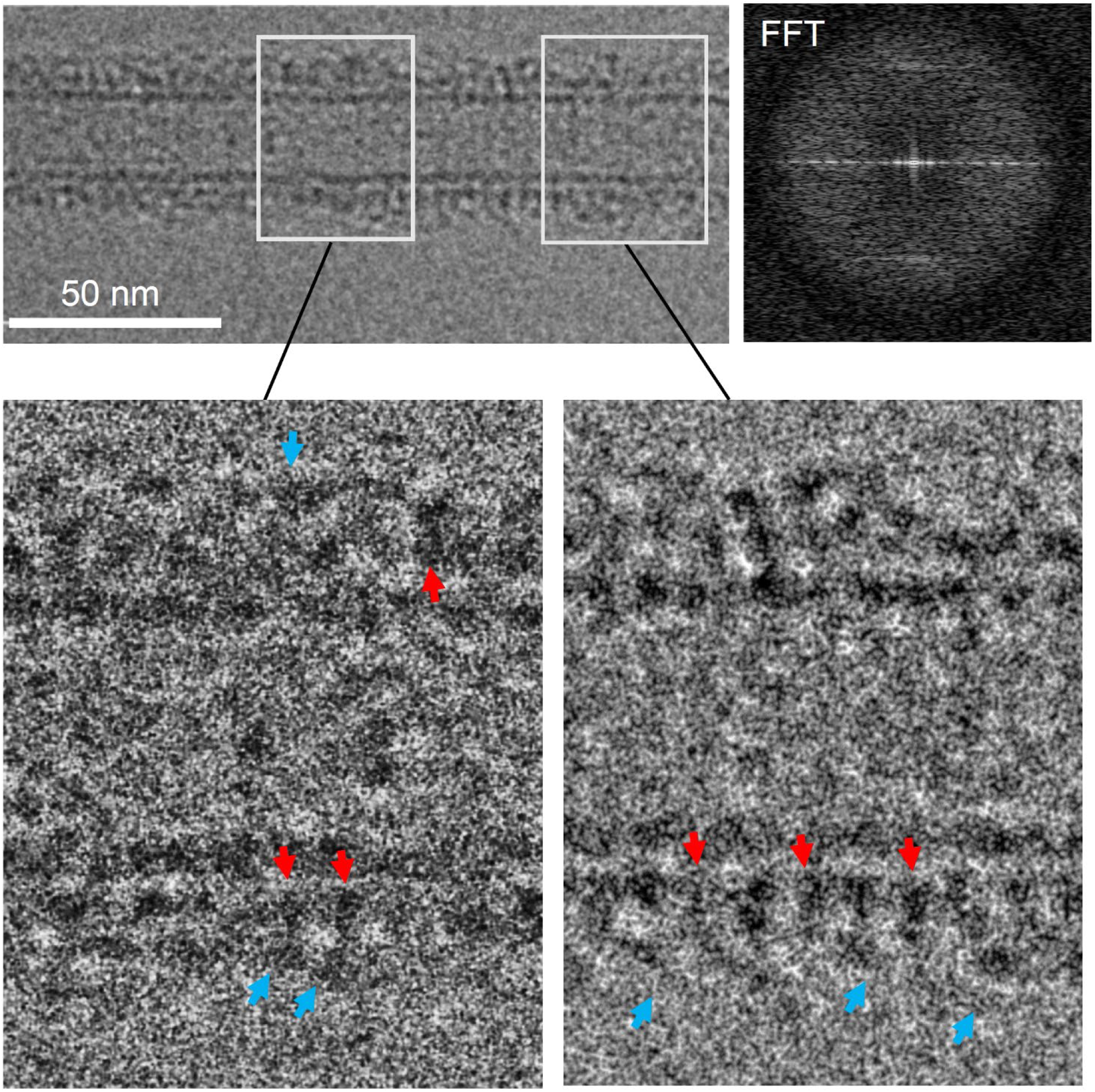
Ca^2+^ disrupts the Syt1^C2AB^-SNARE organization on the lipid membrane. Cryo-EM micrograph of the Syt1^C2AB^-SNARE decorated lipid tube in the presence of 1 mM Ca^2+^ (top left) shows that proteins heavily decorate the LNT surface without forming any pronounced organized structures. Correspondingly, Fourier transform of the vertically oriented tube (top right) shows smeared diffraction peaks. Magnification of the decorated LNTs (bottom) shows regions of both densely (right) and sparsely (left) packed protein on the LNT surface (red arrows), with scattered elongated densities (cyan arrows) extending out from the lipid surface. The protein density on the membrane surface (red arrows) is best approximated by the Syt1 C2 domain, with the SNAREpins corresponding to the auxiliary density (cyan arrow) on the top. It appears that the SNAREpins are displaced from the primary-binding site on the Syt1 protein.

## DISCUSSION

Data presented establish that under resting conditions (1 mM Mg^2+^), Syt1^C2AB^ adopts a previously unknown, molecularly distinct orientation on the membrane surface with partial insertion of the aliphatic loops into the phospholipid membrane. The structure further reveals that the Syt1 C2B domain simultaneously binds the t-SNARE helices on the opposite surface via the recently described ‘primary’ interface and that Syt1 is sandwiched between the SNARE complex and the lipid membrane. In fact, the SNARE complex sitting atop Syt1 is positioned such that its C-terminal end is furthest away from the membrane (Fig. 2). Considering that the SNARE proteins are anchored in the opposing membranes via the C-terminal TMDs, this implies that the Syt1 C2B domain creates a steric constraint to delay complete SNARE assembly by holding the membranes apart. In fact, modeling shows zippering beyond the +5 hydrophobic layer (the middle portion of c-termini) will be sterically impeded due to the separation imposed by Syt1 (Fig. 6). Hence, the Syt1-SNARE configuration on the membrane could inherently contribute to the strength of an overall fusion clamp under resting (1mM Mg^2+^) conditions (Fig. 6). A recent X-ray structure of the Syt1-Cpx-SNARE complex shows that two Syt1 C2B molecules can bind a single SNARE complex - one via the primary site and a second by the Cpx-dependent ‘tripartite’ site^42^. Thus, we speculate that the above described steric clamp might be augmented by an independent C2B domain (derived from Syt1 or Syt7)^44,58^ bound, in conjunction with Cpx, to the to the tripartite interface juxtaposed to the SV membrane (Fig. 6). The resultant ‘dual’ clamp arrangement^44,58^ would trap each SNAREpin in a vice-like grip between the two membranes, held both from above (tripartite C2B on SV) and from below (primary C2B on PM) (Fig. 6).

**Figure 6.**
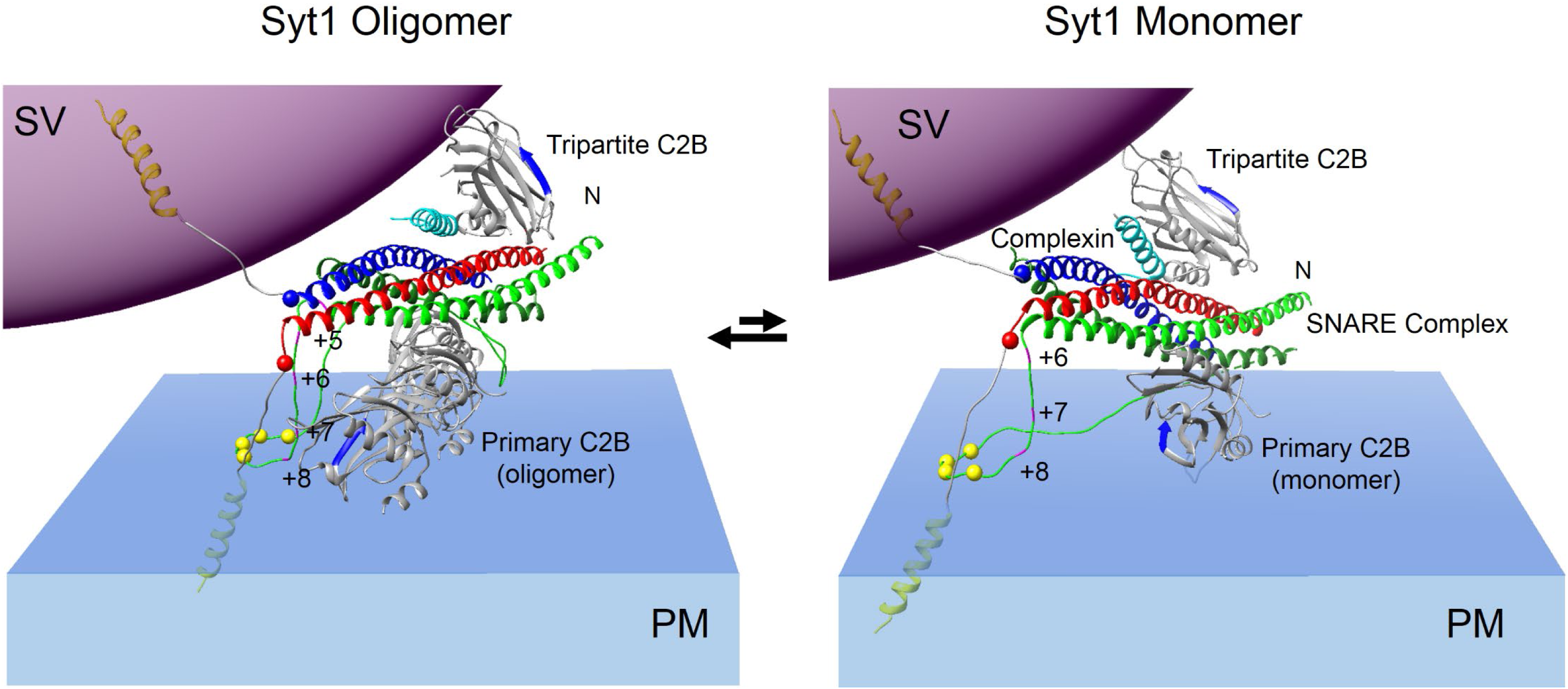
Molecular model for the regulation of SNARE-mediated fusion by Syt1. Upon docking of the synaptic vesicles, Syt1 C2B domains self-assemble into oligomeric structures triggered by PIP2 on the PM, with the aliphatic loops locating to the dimer interface^44,65^. Positioning of the SNAREpin on these Syt1 oligomers via the ‘primary’ binding site would induce steric impediment to the complete zippering of the SNARE complex and in fact, rigid-body fitting shows that zippering beyond layer +4 would be impeded due to the separation imposed by the Syt1 oligomers. Besides the steric block, the oligomers would also radially retain the zippering SNAREpins and balance the assembling forces to generate a stable clamp. This oligomeric clamp may be further augmented by an independent C2B (Syt1 or Syt7) which is positioned to interact with the SV membranes to provide a ‘vice-li’ke dual clamp. Note: Only a part of the ring-like oligomer is shown for clarity. It is possible that these Syt1 oligomers dynamically break and re-form, and thus are not be always intact. Under these conditions, the dissociated Syt1 monomers would re-orient in the membrane corresponding to the Mg^2+^-bound state, with partial membrane insertion of the aliphatic loops, but largely still preserving the SNARE clamped architecture. Thus, Syt1 C2B domains, both as an oligomer or monomer can block full-zippering of the SNARE complex and *vis a vis* fusion under resting conditions. This Syt1-SNARE organization is disrupted following Ca^2+^-influx, as Syt1 reorients into the PM, releasing the attached SNAREpins to fully-assemble and drive fusion.

However, such a clamp is likely to be only meta-stable as the SNAREpins could twist out of position by the radial force generated by SNARE zippering^44^. Additionally, this arrangement cannot account for the observed synchronicity and co-operativity of neurotransmitter release^59-62^. Therefore, while this mechanism may contribute to the overall strength of the fusion clamp, it is unlikely to be the only mechanism at work. So, we envision a concerted mechanism involving the recently described Ca^2+^-sensitive oligomers of Syt1^56,63,64^, which have been shown to be essential to generate a fusion clamp under reconstituted conditions^38^ and for Ca^2+^-control of vesicular exocytosis in PC12 cells^65^. The Syt1 polymerization is driven by the C2B domain, with the Mg^2+^/Ca^2+^-binding aliphatic loops locating to the dimer interface ^56,63,64^. We suspect that under our crystallization conditions, the curved geometry of the LNTs, combined with the C-terminal SNARE interaction stabilizes the membrane-inserted geometry of the Syt1 monomer prevent oligomerization. But under the physiological configuration, with the relatively planar plasma membrane surface and partially-zippered SNARE complex, the Syt1 oligomerization should be more favorable. In fact, Syt1 oligomers are observed on lipid monolayer surfaces even in the presence of 1mM free Mg^2+^ ^56^,^63^.

We posit that Syt1 self-organizes at the site of SV docking triggered by PIP2 clusters on the PM^44,56,63^ and the SNAREpins are positioned on the Syt1 oligomers via the primary interface, similar to the Syt1 monomer (Fig. 6). Indeed, the ‘primary’ binding site is accessible and free to interact with the SNAREpins in the Syt1 oligomer configuration^44,56^. Consequently, the Syt1 oligomer could serve as the template to link multiple SNAREpins to enable rapid and co-operative release^44^. Besides the steric impediment, the Syt1 oligomers will radially restrain the assembling SNAREpins and symmetrically balance the SNARE assembling forces to create a stable clamp on SV fusion^44^. The oligomeric clamp could be further reinforced by the dual clamp arrangement^44,58^ (Fig. 6). Supporting this, recent cryo-electron tomography analysis in PC12 cells revealed a symmetrical organization of the exocytotic machinery under docked vesicles, templated by the Syt1 ring-like oligomers^66^.

It is possible that the C2B oligomers dynamically break and re-form and intact ring-like oligomers are not always present or predominate *in vivo*. Under these conditions, dissociated monomers by re-orienting into the membrane, could still maintain the half-zippered SNAREpins. Thus, Syt1 organization on the membrane, both as a monomer or an oligomer, would lower the probability of fusion of the docked vesicle in the absence of the Ca^2+^-trigger (Fig. 6). Likewise, Cpx is an essential component of this fusion clamp. The Cpx central helix is required to generate the ‘tripartite’ binding site for the second C2B domain^42^ and thus, critical for the ‘dual-clamp’ arrangement^44^ (Fig. 6). Additionally, the Cpx accessory helix has been shown to directly compete with VAMP2 c-terminus zippering and thus, stabilize the half-zippered SNAREpins^67-69^. However, the precise molecular arrangement that enables the accessory-helix based clamp is still unclear.

The Syt1-SNARE organization on the LNT surface is disrupted upon Ca^2+^ addition (Fig. 5). It appears that as the Syt1 C2 domains reorient into the membrane following Ca^2+^ binding, the associated SNARE complexes are displaced from the primary binding site. Modeling suggests that Syt1 reorientation into the membrane would produce a steric clash between the SNARE complex bound to the primary site and the vesicle/plasma membranes (Supplementary Fig. 9). Previous fluorescence analyses suggest that some form of Syt1-SNARE interaction is maintained during the Ca^2+^-activation process^70,71^. Given the low signal-to-noise ratio of the EM map, we cannot rule out this possibility, but the precise nature of this interaction remains unclear. Nonetheless, our data argues against the *en bloc* activation model^41,47,70-72^ and suggests that a significant conformational rearrangement is associated with the Ca^2+^-activation process. Incidentally, the Ca^2+^-binding also reorients the Syt1 from the ring oligomer geometry, disrupting the oligomers^56,63^. Thus, it emerges that the Ca^2+^-activated conformation changes in the Syt1 C2B domain, which is physiologically required for triggering synaptic transmission^21,24,25^, concomitantly reverses the Syt1 clamp and liberates the partially assembled SNAREpins to complete zippering and open the fusion pore cooperatively. Although speculative, the above described model provides a plausible framework to explain the kinetics of Ca^2+^-triggered SV fusion in molecular terms. Additional focused high-resolution structural and functional analysis is required to ascertain its relevance.

In summary, our data shows that the divalent cation (Mg^2+^/Ca^2+^) coupled membrane interaction is a key element of Syt1 function as both a clamp and an activator of SNARE-mediated fusion. Under resting conditions (with Mg^2+^), Syt1 locates between the SNARE complex and the plasma membrane creating a SNARE complex-specific steric constraint to prevent full-zippering (Fig. 6). Ca^2+^-triggered reorientation into the lipid membrane allows the associated SNAREpins to complete zippering and trigger fusion. In this manner, Syt1 couples Ca^2+^-influx to SNARE-mediated SV fusion and neurotransmitter release.

## MATERIALS AND METHODS

Lipids, 1,2-dioleoyl-sn-glycero-3-phosphocholine (DOPC), 1,2-dioleoyl-sn-glycero-3-phospho-L-serine (sodium salt) (DOPS), L-α-phosphatidylinositol-4,5-bisphosphate (Brain, Porcine) (ammonium salt) (PIP2), D-galactosyl-ß-1,1’ N-nervonoyl-D-erythro-sphingosine (GC) were purchased from Avanti Polar lipids (Alabaster, AL). Thiol reactive fluorescent probe IAEDANs (1,5-IAEDANS, 5-((((2-iodoacetyl)amino)ethyl)amino)naphthalene-1-sulfonic Acid) was purchased from ThermoFisher Scientific, Waltham, MA. The Syt1^C2AB^-SNARE construct^41^ was generously provided by Dr. Axel Brunger (Stanford University). This included two duet plasmid constructs: One containing both the Syt1 C2AB domain (residue 141-421) covalently linked to the N-terminus rat SNAP-25 (SNAP25 SN2, residue 141-204) by a 37 amino acid linker, and the N-terminus of SNAP-25 (SNAP25, SN2, residue 7-83). This is co-expressed with a second plasmid containing the rat syntaxin-1A fragment (residue 191-256), and the His-tagged rat synaptobrevin-2 SNARE domain (residue 28–89). The fluorescence analyses were carried out using rat synaptotagmin-1 C2AB domains (residues 143-421). Single cysteine mutants (M173C, S235C, V304C and I367C) were generated using a QuickChange mutagenesis kit (Agilent Technologies).

### Protein Expression and Purification

The Syt1-SNARE complex was purified as described previously^41^ with minor modifications. Briefly, E. coli BL21 (DE3) was grown to an OD_600_ of 0.6-0.8, induced with 0.3 mM isopropyl ß-D-1-thiogalactopyranoside (IPTG), the cells were harvested after 4h at 37°C and placed in −80°C until purification. The pellets were suspended in lysis buffer (50mM Tris-HCl, pH 8.0, 350mM KCl, MgCl_2_ 1mM, CaCl_2_, 20mM imidazole, 0.5mM TCEP, 10% Glycerol) with addition 1% Triton X-100 and EDTA-free protease inhibitor cocktail (Sigma) and lysed by a cell disrupter. Before the centrifugation (100,000xg for 30 min), the lysate was supplemented with 0.1% polyethylemine (pH 8.0). The supernatant was loaded onto Ni-NTA beads (Thermo-Scientific) equilibrated in the lysis buffer with 0.1% Triton X-100. Beads were harvested and washed with lysis buffer with 0.1% Triton X-100 and incubated with the lysis buffer supplemented with 10 mg/ml DNAase and 10 mg/ml RNAase and at room temperature for 45 min. The beads were washed with lysis buffer supplemented with 1mM CaCl_2_ and 30 mM Imidazole. The Syt1–SNARE complex was eluted in the lysis buffer supplemented with 330 mM imidazole. The His-tag was cleaved overnight using TEV protease during dialysis (dialysis buffer: 50mM Tris-HCl, pH 8.0, 100mM KCl, 1 mM CaCl_2_, 0.5mM TCEP, 10% Glycerol) overnight at 4C°. The un-cleaved proteins were removed with Ni-NTA beads and then subjected to anion exchange chromatography (buffer A: 50 mM Tris-HCl pH 8.0, 50mM KCl, 1 mM CaCl_2_, 0.5mM TCEP, 10% Glycerol; buffer B: 50mM Tris-HCl, pH 8.0, 1M KCl, 1 mM CaCl_2_, 0.5mM TCEP, 10% Glycerol) using a linear gradient of KCl starting at 50mM and ending at 1 M. The pooled peak fractions (KCl concentration ∼240 mM) were concentrated, flash-frozen in liquid N_2_ and stored at −80°C until further usage. The quality of the protein was assessed by SDS-PAGE. Only the protein sample that demonstrated stability and efficacy when comparing un-boiled samples versus boiled was used in experiments. The Syt1^C2AB^ construct (aa 143-421) with cysteine modifications were expressed and purified as described previously^56,63^.

### Model membranes and Lipid Nanotube preparation

Vesicles were prepared with lipid composition of DOPC/DOPS/PIP2 in molar ratio 60/34/6. The lipid stocks were mixed together in chloroform and methanol mix and the solvent was evaporated under N_2_ and the samples were vacuum desiccated for 1 hour. The resulting dried lipid film was hydrated for 1 hour at room temperature with constant vortexing in the appropriate buffer. For structural studies, the buffer composition was 20 mM MOPS pH 7.4, 5 mM KCl, 1 mM EDTA, 0.5 mM TCEP, 250 uM EGTA; for fluorescent-based membrane penetration assays, the labeling buffer was used with 0.2 mM EDTA. Liposomes were prepared using multiple freeze-thaw cycles and then extruded (Mini-Extruder, Avanti Polar Lipids) through a 400 nm pore filter (Whatman) for structural studies and a 50 nm pore filter for fluorescence experiments.

To prepare lipid nanotubes (LNT), Galactyl ceramide (GalCer 24:1) was included in the lipid composition. Two different compositions of LNTs were used: DOPC/GC/DOPS/PIP2 and GC/DOPS with lipid molar ratio 40/20/34/6 and 20/80, respectively. The lipids were mixed in chloroform, dried and then hydrated using 20 mM MOPS pH 7.4, 5 mM KCl, 1 mM EDTA, 0.5 mM TCEP buffer. The samples were sonicated using a bath sonicator (Branson Ultrasonics, Danbury, CT) and self-assembled LNTs were stored at 4C.

### Helical crystallization of Syt1^C2AB^-SNARE complex

Purified Syt1^C2AB^-SNARE complexes were made up to a stock concentration to ∼40 mM so that the final protein and salt concentration (∼25 mM KCl) were similar under all experimental conditions. Protein dilution was performed in low-salt buffer (20 mM MOPS pH 7.4, 5 mM KCl, 1 mM EDTA, 0.25 mM EGTA, 2% (w/v) Trehalose (Sigma), 0.5 mM TCEP and mixture of MgCl2 or CaCl2 to provide a 1mM final concentration of free ions (based on MaxChelator online calculator). For the liposome samples, pre-formed liposomes (∼0.5 mM lipids) were applied to glow-discharged homemade carbon coated copper grids (400 mesh). After 20 min incubation at room temperature in a humidity box, the excess vesicles were removed by washing and a 4 μM protein solution was added to the grids. The samples were either negatively stained or frozen for Cryo-EM after 1hour incubation at room temperature in a humidity box. For the LNT samples, LNT suspension was mixed with the protein using 0.5 mg/ml lipids + 0.5 mg/ml protein in 1:1 volume ratio. The final protein and KCl concentration were ∼4 μM and ∼30 mM respectively. After 30 min incubation at room temperature samples were negatively stained or flash-frozen on Vitrobot Mark III. Helical crystals were observed at 100 mM KCl and confirmed by negative staining microscopy, but those samples showed high sensitivity to the freezing process since most of the tubes in the Cryo-EM samples were naked.

### Fluorescent Labeling

Syt1^C2AB^ constructs with single cysteine residues (M173C, S235C, V304C and I367C) were mixed and incubated with a 10-fold molar excess of the IAEDANS dye (freshly prepared) in HEPES buffer (40 mM HEPES pH 7.4, 100 mM KCl, 0.5 mM TCEP, 10% Glycerol) at 4°C overnight. Labeled proteins were separated from the free fluorophore using a NAP-5 Sephadex G-25 desalting columns in labeling buffer (GE Healthcare). The labeling efficiency was determined using an extinction coefficient of 5400 M^-1^cm^-1^ at 335 nm for IAEDANS and the protein concentration was determined by Bradford assay using BSA as a standard. Labeling efficiency was >85% in all cases.

### Fluorescence Measurements

Labeled protein was mixed with liposomes in 1:1000 molar ratio (0.5 μM protein: 0.5 mM lipids) and incubated at 4°C overnight to allow for complete binding. Prior to the fluorescence measurements, the samples were brought up to room temperature by incubating 30 min in a 25°C water bath. Steady-state fluorescence measurements were made at 20°C using a PC1 photon counting spectrofluorimeter (ISS, Illinois). IAEDANS probe was excited at 336 nm, and emission spectra were collected from 400 to 600 nm using a quartz cuvette with a 2-nm slit. Resulted emission spectra were corrected using empty vesicles (without protein) prior to analysis.

### Electron microscopy sample preparation and data acquisition

Negative staining of samples was done using home-made carbon coated copper grids (400 mesh). Vesicles samples were stained right after the incubation time, lipid nanotubes samples were applied onto glow-discharged grids for 1 min. Both samples were stained with uranyl acetate solution (1% w/v) and air dried. Negatively stained samples were imaged using FEI Tecnai T12 operated at an acceleration voltage of 120kV, equipped with LaB6. Micrographs were recorded using 4k x 4k Gatan UltraScan 4000 CCD camera with nominal magnification of 42000x and under low-dose conditions (∼25e^-^/Å) with defocus range of 1-2.5 um. Micrographs were binned by a factor 2 at a final sampling of 5.6 Å per pixel.

For the Syt1^C2AB^-SNARE decorated vesicles, the samples on the grids were pre-washed with incubation buffer without trehalose, blotted with paper while instantaneously adding 2.5 μl of the same buffer, transferred into a Vitrobot Mark III held at 20°C with 100% humidity, blotted for ∼5sec and plunge-frozen in liquid ethane cooled by liquid nitrogen. For the Syt1^C2AB^-SNARE decorated lipid nanotubes, the C-flat holey carbon grids (Protochips, CF-1.2/1.3-3C-T) were plasma cleaned with Solarus Model 950 plasma cleaning system (Gatan) using O_2_/Ar mixture for 15 sec. Then 2.5 ul of the sample was applied to the grids, blotted for ∼7 sec and plunge-frozen in liquid ethane cooled by liquid nitrogen using a Vitrobot Mark III at 20°C with 100% humidity.

Images were recorded on FEG equipped FEI Tecnai F20 microscope operating at 200 kV at a nominal magnification of 25,000 on Gatan K2 Summit direct detection camera in counting mode with resulted pixel size 1.485 Å. Each image was dose-fractionated to 33 frames with a total exposure time 9.9 sec, 0.3 sec per frame. The total dose was ∼44 e^-^/ Å^2^. Movies were recorded using SerialEM^73^with nominal defocus values from −1.5 to −3.0 um.

### Cryo-EM data processing

Movies were motion-corrected, dose-weighted and summed using Unblur^74^. Micrographs were evaluated for presence of helically organized tubes by visual appearance and helical diffraction pattern of FFT using DigitalMicrograph (Gatan) and Fiji (ImageJ). Helically organized tubes were sorted by outer diameter measured manually in DigitalMicrograph. Segments of best representatives from different diameter groups with clear layer lines in reciprocal space were selected for indexing with EMIP^52^. Surprisingly, only small population of the tubes with outer diameter ∼41nm showed helical diffraction pattern corresponding to homogenous helical crystal and was suitable for obtaining real-space helical parameters as rise (Δz) and twist (Δφ). Calculated possible parameters of helical lattice were checked by low resolution IHRSR reconstruction using SPARX^75^. Briefly, using *boxer* from EMAN1 package^76^ implemented in SPARX particles were extracted with 97% overlap and three times binned and used in SPARX-IHRSR reconstruction using featureless hollow cylinder created using SPIDER^77^. Parameters that gave reconstruction with pronounced, elongated densities with similar dimensions as SNARE-pins (rise, Δz=7.35 Å and twist, Δφ=78.48°; 14,-4 start helix) were used for further high-resolution reconstruction in RELION 2.0^78,79^. For the final reconstruction all tubes with an outside diameter approximately 41 nm (as identified above) were manually selected using *e2helixboxer.py* from EMAN2^80^ and boxes were extracted with an inter-box distance of 3 asymmetrical units (96% overlap) using RELION 2.0. Defocus values were estimated by CTFFIND4^81^. The resulting 2844 particles were subjected to 2D classification and particles belonging to blurry classes were discarded, leaving 2082 particles that were further subjected to 3D-refinement using a featureless hollow cylinder as an initial reference. The resulting 3D volume was post-processed using soft mask and applying a B-factor of −300. The average resolution estimated using a gold standard (FSC=0.143) was 10.4 Å (Supplementary Fig. 10). The local protein resolution estimated using *blocres* program (BSOFT^82^) was in the range of 8-12 Å (Supplementary Fig. 10)., Only one handedness gave density consistent with the X-ray structures of SNAREs, Synaptotagmin C2A and C2B domains. The selected handedness defines a left-handed helix on the tube’s surface (Δφ=-78.48°, Δz=7.35 Å).

### 3D model refinement and model building

First fitting was performed using separate crystal structures of C2A, C2B and the SNARE-pin obtained from a Mg^2+^ bound crystal structure (PDB code: 5CCI^41^) into the Cryo-EM map manually using UCSF Chimera^83^ to help segment the map end extract Cryo-EM densities of several complexes on the surface of outer leaflet. The extracted map was used for rigid-body fitting using SITUS^84^ and top-scoring solutions were selected for the further analysis. All model building was performed using UCSF Chimera.

### Data availability

The 3D CryoEM density map of the Syt1^C2AB^-SNARE decorated LNTs (flipped along z-axis, representing left-handed helix) has been deposited in the Electron Microscopy Data Bank under accession number EMD-9231. The deposition includes the corresponding EM map, both half maps and the mask used for the final FSC calculation. Coordinates for the Syt1^C2A^, Syt1^C2B^, SNAREpins fitted into the map by SITUS have been deposited in the Protein Data Bank under accession number PDB 6MTI. All other data are available from the corresponding authors upon request.

## ACKNOWLEDGEMENTS

This work was supported by National Institute of Health (NIH) grant DK027044 to JER. We are grateful to the staff of the Electron Microscopy & Cryo Electron Microscopy Facility at the Center Cellular and Molecular Imaging, Yale School of Medicine and of the High-Performance Computing Facility for expert support and guidance. We also wish to thank members of the Sindelar lab, particularly Dr. Xueqi Liu, for help with data processing.

