## Supplementary Information for "Structural Basis for the Clamping and Ca^2+^ Activation of SNARE-mediated Fusion by Synaptotagmin"

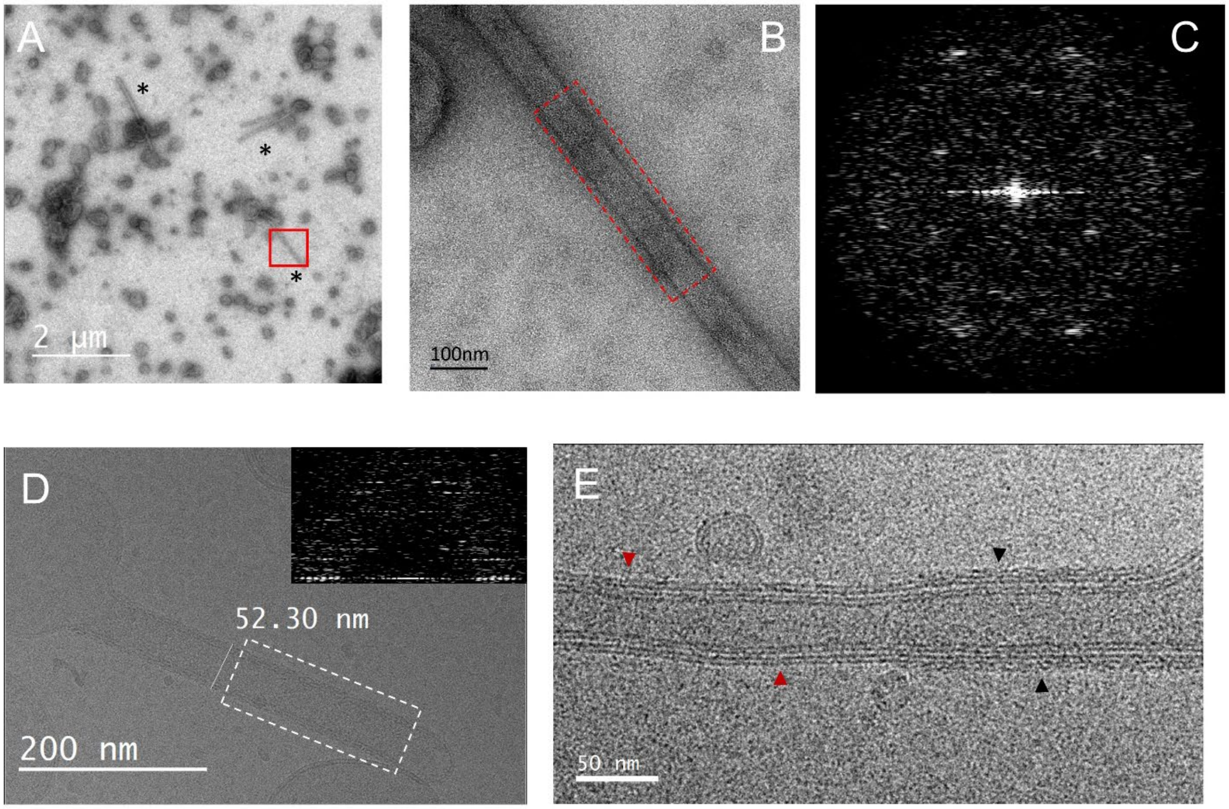


**Supplementary Figure 1**. Syt1$^{C2AB}$-SNARE complex forms helically organized tubes out the negatively charged vesicles (DOPC/DOPS/PIP2 60/34/6). (A) Negative staining microscopy of vesicles on a carbon surface after 1-hour incubation at room temperature with Syt1$^{C2AB}$-SNARE protein in the presence of 1 mM $Mg^{2+}$. Only few vesicles are transformed into tubes (black asterisk) by the protein. (B) Magnified view of the tube highlighted by the red box in A. (C) Fourier transform of the selected tube area in B. with diffraction peaks corresponding to helically organized protein on the tube's surface. (D) Cryo-EM micrograph of a tube formed by Syt1$^{C2AB}$-SNARE complex. The fourier transform of the highlighted area is shown in the inset. Unfortunately, these well-diffracting tubes were a minor fraction, while most of the tubes had severe flaws in the protein coating (E). Most of the tubes lacked protein coverage (red triangle) or were only partially decorated (black triangle), probably due to damage during the vitrification process.

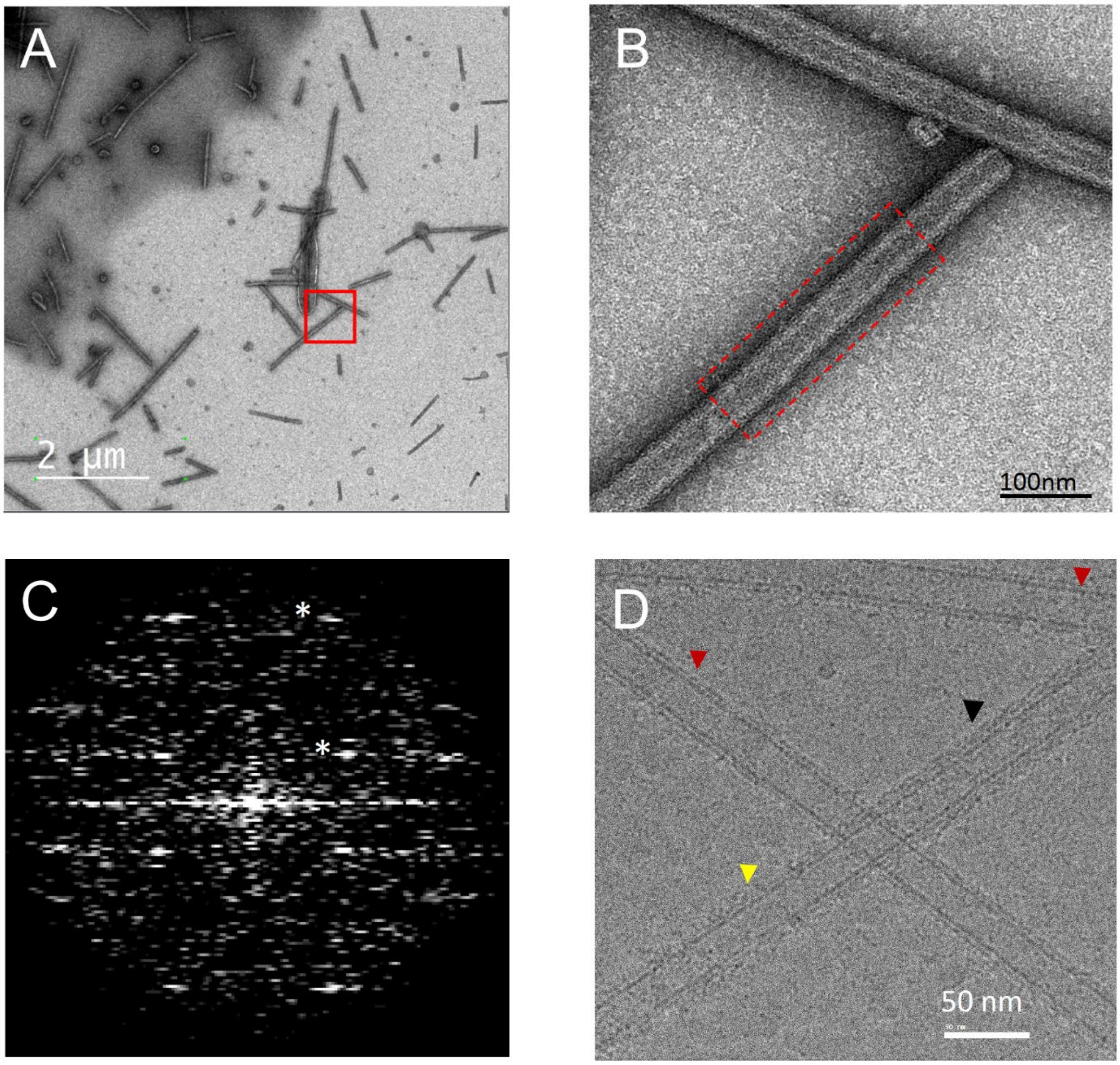


**Supplementary Figure 2.** Syt1$^{C2AB}$-SNARE complex helically organizes on preformed negatively charged lipid nanotubes (DOPC/GalCer/DOPS/PIP2 40/20/34/6). (A) Negative staining microscopy of lipid nanotubes after 30 min incubation at room temperature with Syt1$^{C2AB}$-SNARE protein in the presence of 1mM $Mg^{2+}$. (B) Magnified view of the selected area in (A). The tubes showed a regularity in protein coating which was confirmed by Fourier transform (C) of the selected tube area with diffraction peaks corresponding to helically organized protein on the tube's surface. Main peaks pattern (white asterisk) is similar to vesicular tubes diffraction (Fig. S1C). (D) Similar to the vesicle samples, Cryo-EM micrograph of LNTs sample showed a majority of empty tubes (red triangles) or tubes without helically organized protein (yellow triangle), with small regions of helically organized protein (black triangles). This indicated that the Cryo-EM sample preparation process damages the sample.

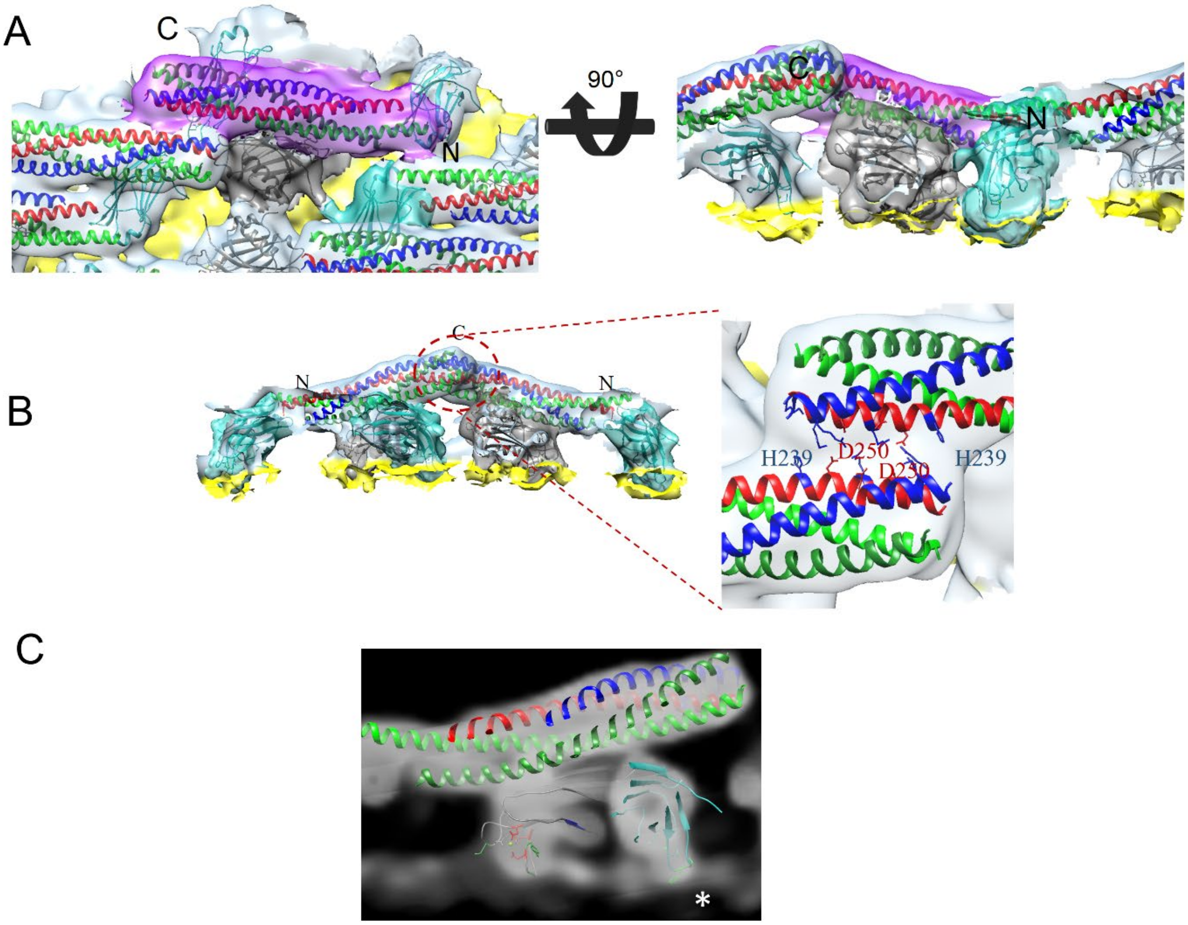


**Supplementary Figure 3.** (A) 3D-surface representation of a single unit of Syt1$^{C2AB}$-SNARE complex (PDB code: 5CCI) consisting of the primary interface alone fitted into the helical crystal on the membrane surface. Both the top view (left) and side-view (right) are shown. The SNAREpin (magenta) is inclined with its C-terminus oriented outward from the membrane, while the C2B (gray) and C2A (blue) domains are positioned between SNAREs and the membrane. Both domains interact with top layer of the tube which corresponds to phospholipid heads plane of the outer leaflet, with a visible penetration of the aliphatic loops into the membrane surface. (B) The 2D helical structure is stabilized by interaction between neighboring SNAREpins. This involves polar interactions between charged residues in the C-terminus of the neighboring Syntaxin-1 (red) and VAMP2 (blue) helices. (C) Density slice through the C2B (gray ribbon) and C2A (blue ribbon) clearly showing protein density penetration into the membrane phospholipid heads plane (white asterisk).

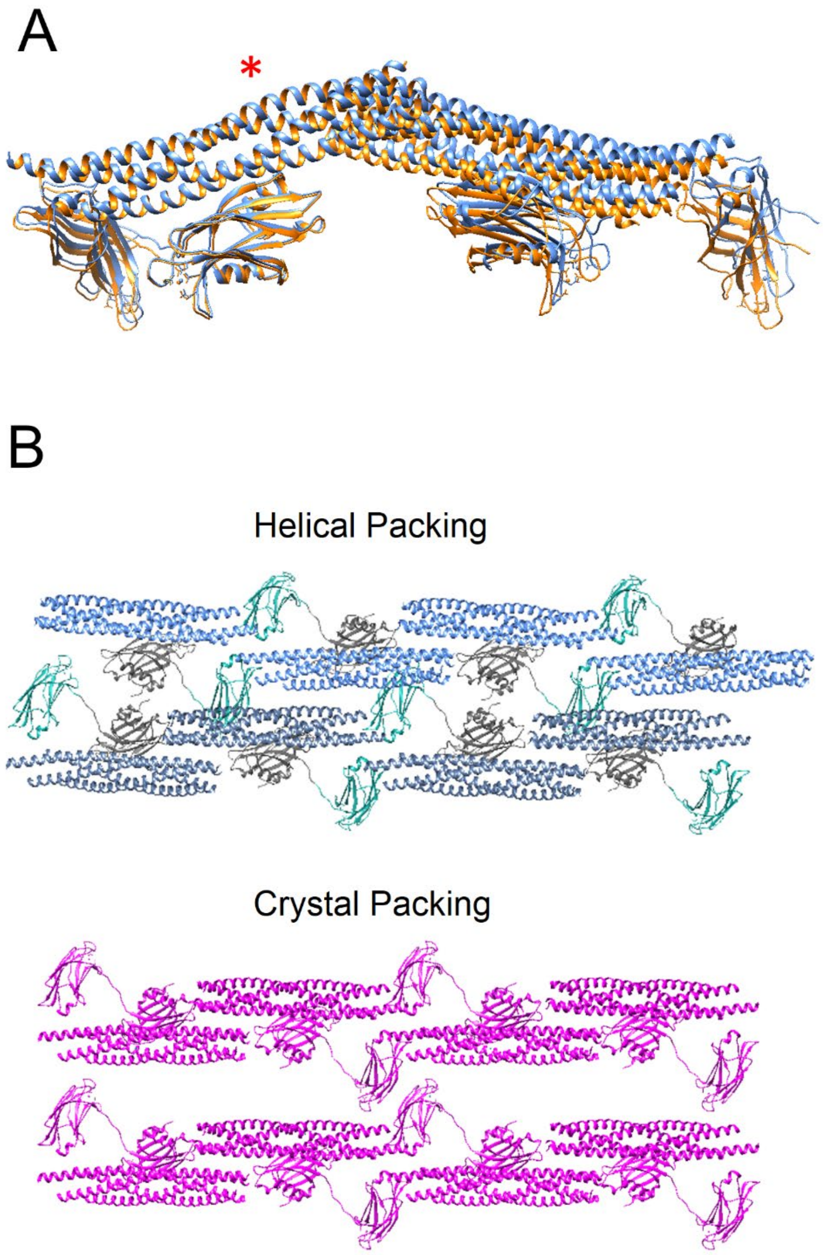


**Supplementary Figure 4.** Comparison of helical packing on the membrane surface and 3D crystal formed in the presence of $Mg^{2+}$ (PDB 5CCI). (A) Superimposition of a dimer of $Syt1^{C2AB}$-SNARE reconstruction (orange) and 3D-crystal (primary interface unit, blue). Dimers were aligned together using the SNARE domains (red asterisk). C2B domains of aligned SNAREpins have almost a perfect match whereas C2A domain and the adjacent $Syt1^{C2AB}$-SNARE complex show a slight deviation from 3D crystal packing. (B) Top view of the 2D planes from the helical packing (top) and 3D crystal (bottom). $Syt1^{C2AB}$-SNARE complexes are more densely packed on the membrane compared to the plane in the 3D crystal.

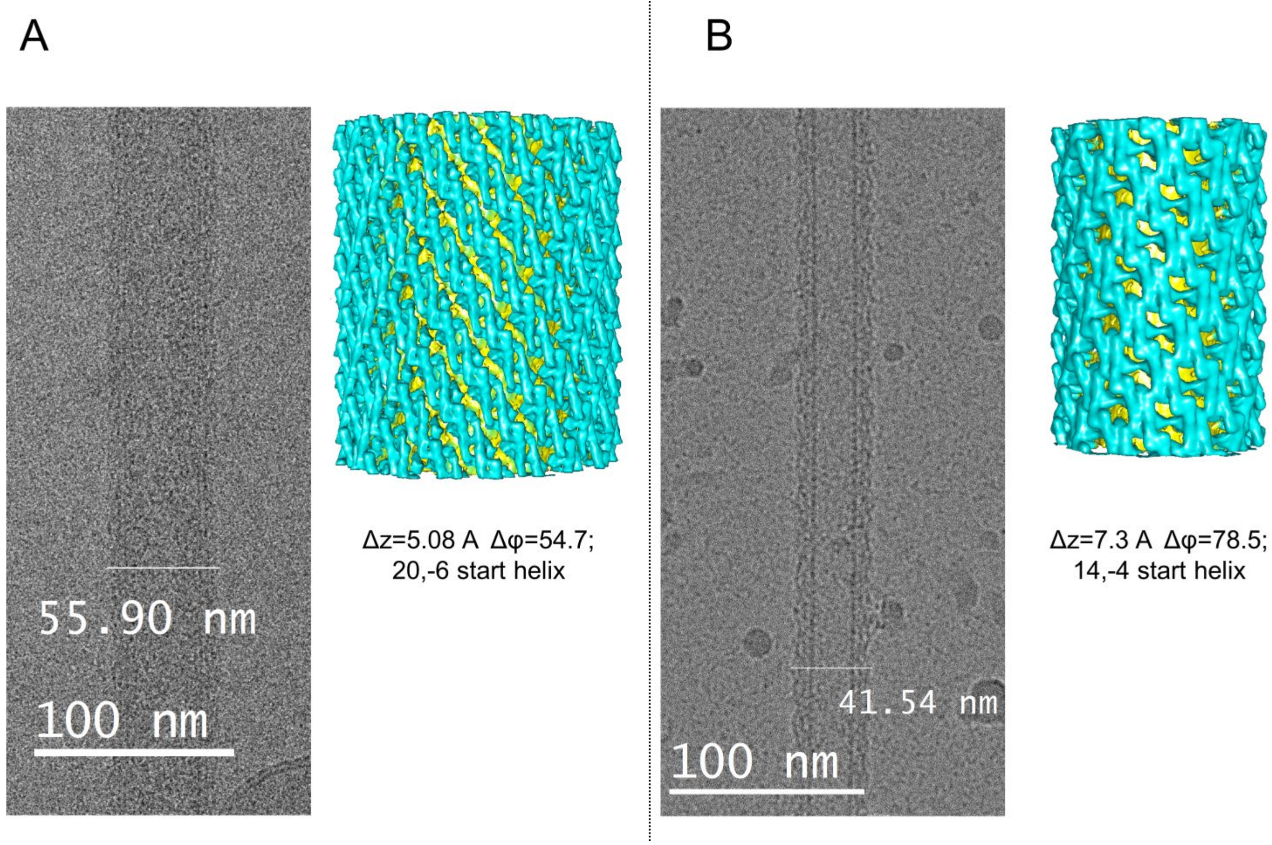


**Supplemental Figure 5.** (A) Cryo-EM micrograph of a representative tube formed from negatively charged vesicle (DOPC/DOPS/PIP2 60/34/6) using Syt1$^{C2AB}$-SNARE protein (left). This single tube was indexed with protein organized as 20,-6 start helix with helical rise $\Delta z$ = 5.08Å and twist $\Delta\varphi$ = 54.7°. The resulted 3D volume after reconstruction in SPARX is shown on the right panel. Despite the low-resolution, it is clear that protein organization on the tube surface is identical to that obtained using 80:20 DOPS:GC LNTs (Figure 1) (B) Representative cryo-EM micrograph of Syt1$^{C2AB}$-SNARE protein helically organized on lipid nanotubes (left panel) with a lipid composition close to that used for the vesicles (DOPC/GC/DOPS/PIP2 40/20/34/6). The indexing showed that tubes with identical dimensions have the same helical parameters (14,-4 start helix, $\Delta z$=7.3; $\Delta\varphi$=78.5;) and the resulting low resolution 3D volume (right) from SPARX is similar to the high-resolution reconstruction (Figure 1).

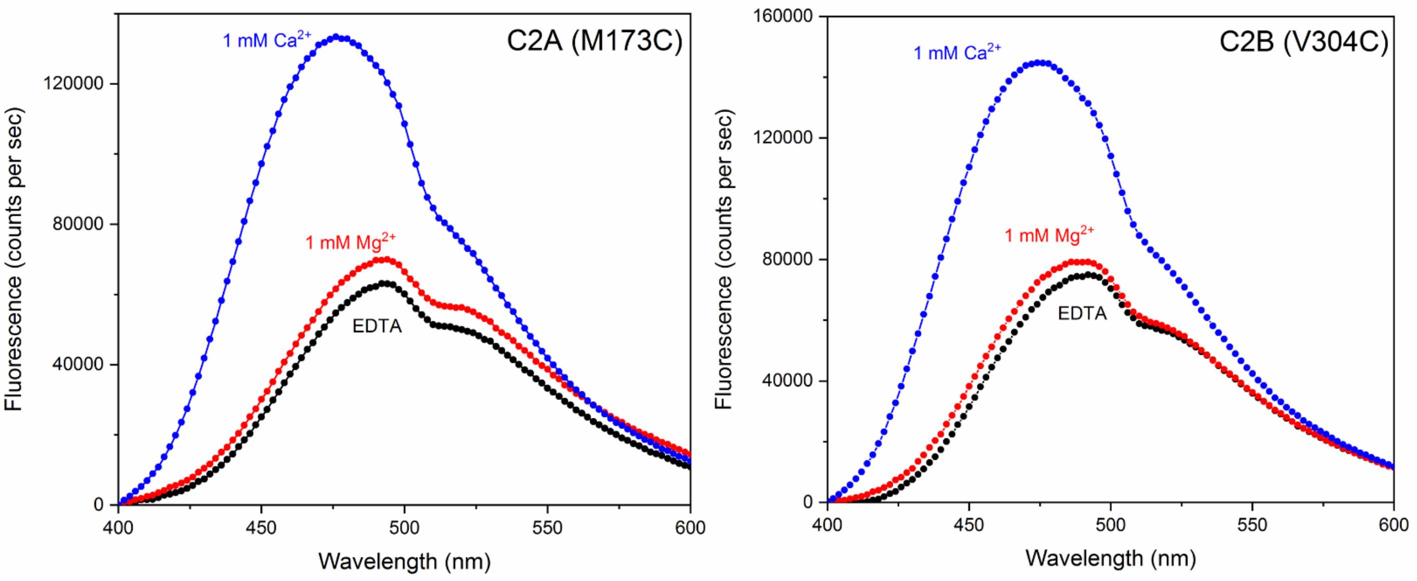


**Supplementary Figure 6**. Representative fluorescence emission curves of environmentally-sensitive probe, IAEDANS introduced at the tip of the aliphatic loop 1 of the C2 domains (residue M173 on C2A & residue V304 on C2B) showing loop 1 inserts into the phospholipid bilayer only following $Ca^{2+}$-influx. Accordingly, we observed no change in the fluorescence intensity in the presence of $Mg^{2+}$ (red curve) as compared to the EDTA control (black curve), whereas $Ca^{2+}$ addition (blue curve) resulted in a large increase in the fluorescence intensity and blue-shift in the emission maxima, corresponding to membrane localization of loop 1 residues. This is in contrast to the loop 3 behavior, which is partially inserted with $Mg^{2+}$ and deeply inserted in the $Ca^{2+}$-bound state (Figure 4).

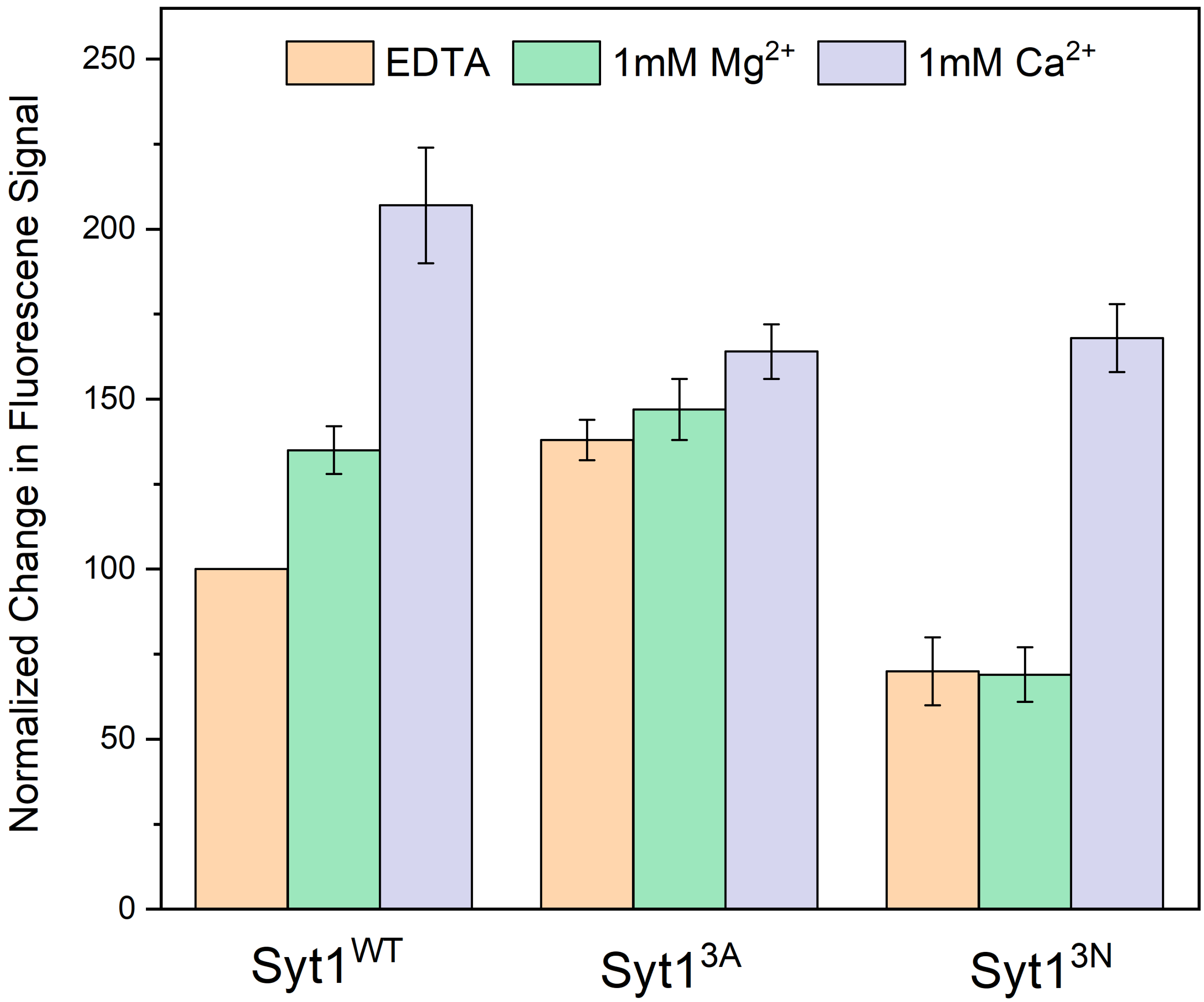


**Supplementary Figure 7.** IAEDANS fluorescence signal introduced in loop 3 of C2B domain (residue I367) was used to determine the effect of altering the physiochemical properties of the $Ca^{2+}$-loops on its membrane-insertion ability. Enhancing the hydrophobicity of the $Ca^{2+}$-loops (D307/ D363/D365A; $Syt1^{3A}$) created a constitutively partially membrane-inserted state, which formed even in the absence of $Mg^{2+}$ and was not disrupted by $Ca^{2+}$ addition. In contrast, polar mutations in the loop (V304N/Y364N/G368N; $Syt1^{3N}$) abolished membrane insertion under EDTA and $Mg^{2+}$-conditions, but $Ca^{2+}$ binding triggered partial membrane-insertion. This data, taken together with the observation that $Syt1^{3A}$-SNARE complex forms 2D crystals on LNTs under all conditions, but $Syt1^{3N}$-SNARE complex only in the presence of $Ca^{2+}$ (Supplemental Fig. 8), strongly argues that the partially membrane inserted state guides the observed helical assembly on LNT surface.

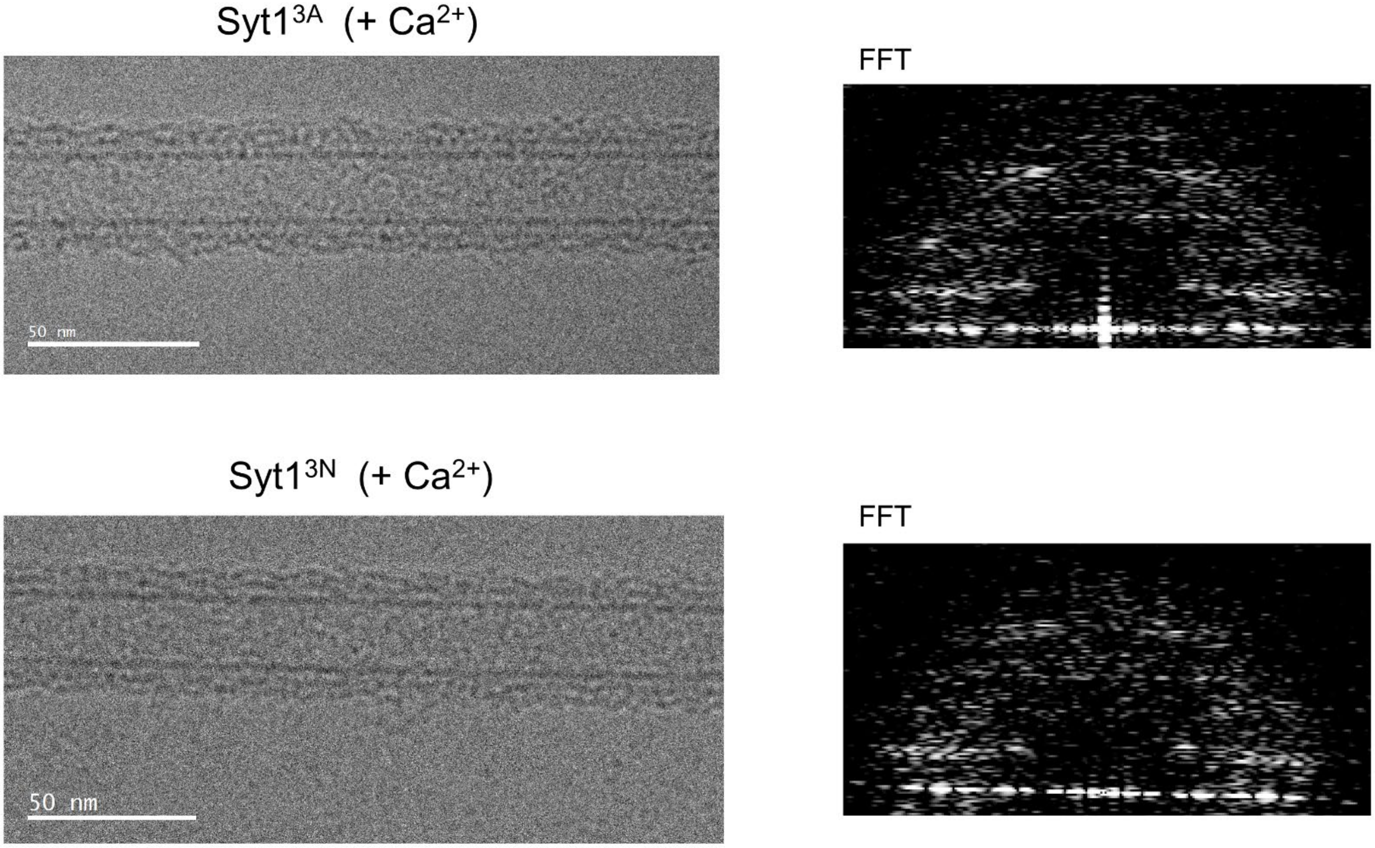


**Supplementary Figure 8.** Helical organization of Syt1$^{C2AB}$-SNARE complex mutants in the presence of $Ca^{2+}$. Top: Unlike Syt1 wild type, the Syt1$^{C2AB}$-SNARE organization on the LNT surface is largely preserved in the Syt1 mutant with compromised $Ca^{2+}$-binding ability (D309A/D363A/D365A, Syt1$^{3A}$) following $Ca^{2+}$-addition. The cryo-EM micrograph of a representative tube area is shown in the left and corresponding Fourier transform of the area is shown on the right. Bottom: $Ca^{2+}$ largely restores the helical order of the Syt1 mutant with decreased hydrophobicity of C2B domain $Ca^{2+}$ loops (V304N/Y364N/I367N, Syt1$^{3N}$) which failed to form a helically organized structure on the LNT surface in the presence of $Mg^{2+}$ (Figure 3). In both cases, we observe small defects in the crystal and smearing of FFT compared to the wild type (Figure 3), likely due to the re-orientation of the C2A domain, which is not affected by the C2B mutations.

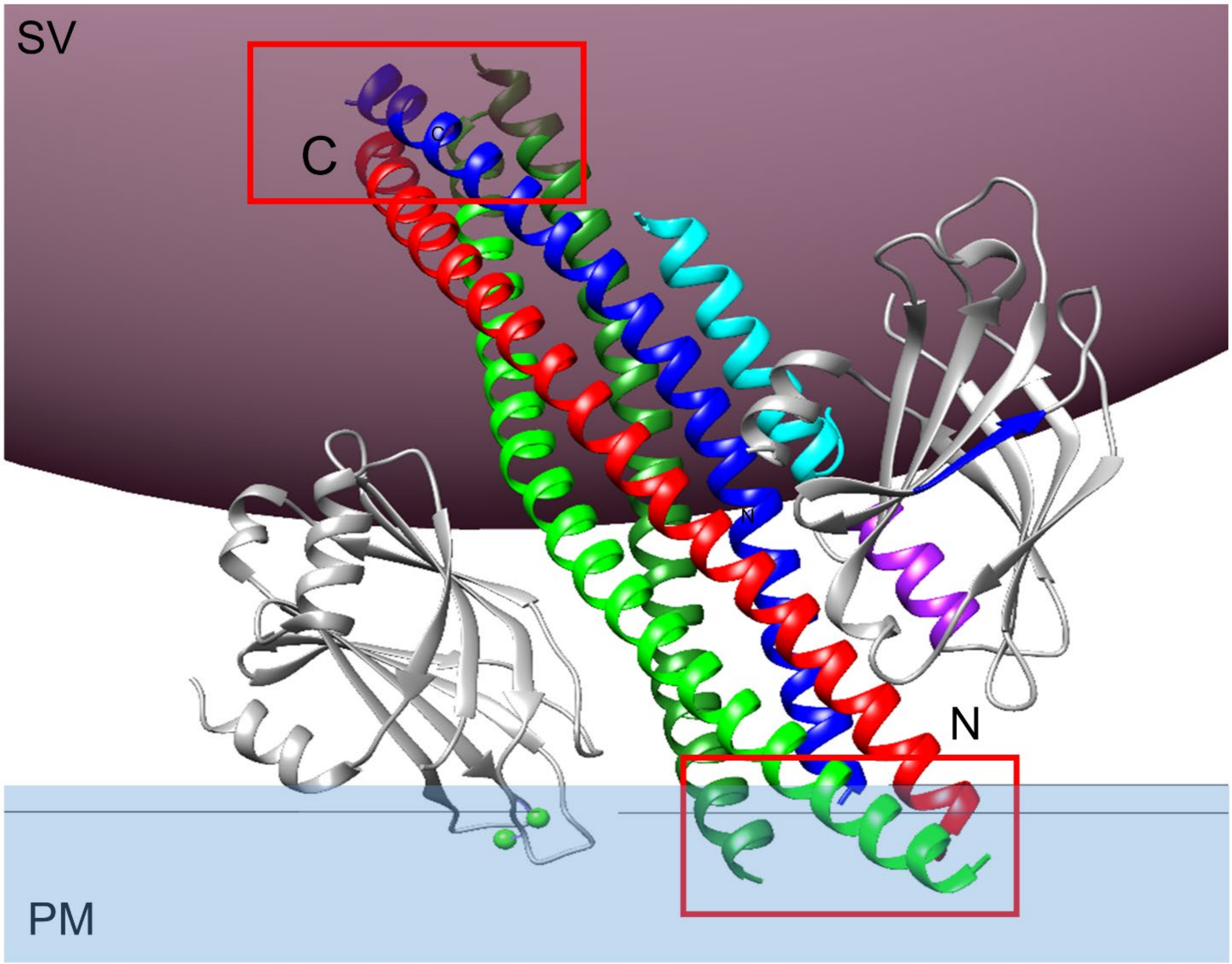


**Supplementary Figure 9**. Modeling shows that the $Ca^{2+}$-induced re-orientation of the Syt1 C2B domains in the lipid bilayer will introduce a large steric clash (red boxes) at both the N- and C-terminal end of the SNARE complex, which are bound to the primary interface and the opposing membranes respectively. This provides a likely explanation for the observed dissociation of the SNARE from $Ca^{2+}$ bound Syt1 (Figure 5).

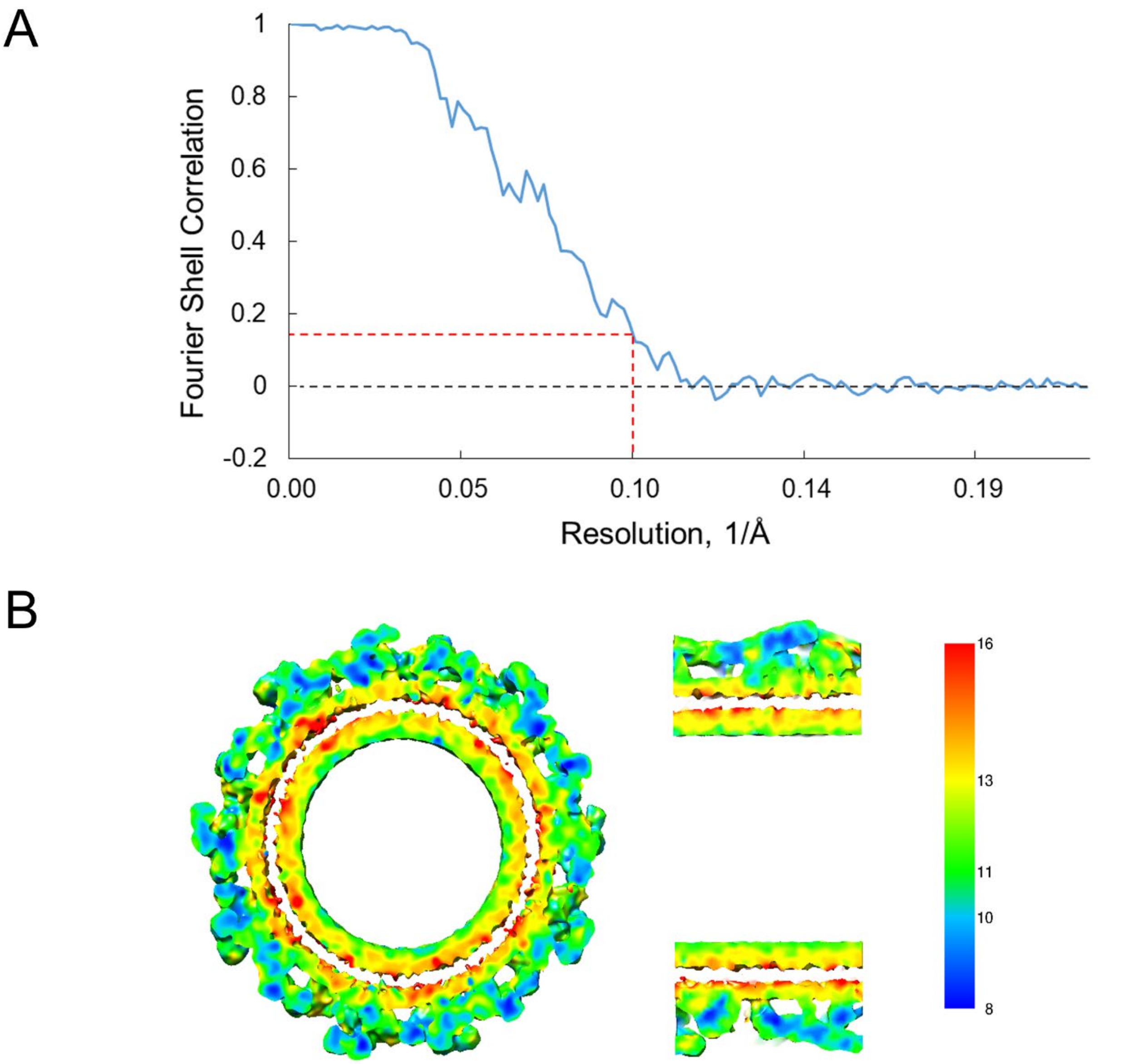


**Supplementary Figure 10**. (A) Gold standard Fourier Shell Correlation (FSC) curve of the Syt1$^{C2AB}$-SNARE complex 3D map. (B) Cross-section of the final Cryo-EM map colored by local resolution as estimated by *blocres* program[1]
